# Shared and organ-specific gene expression programs of fibrotic diseases

**DOI:** 10.64898/2026.03.09.709232

**Authors:** Leonie Küchenhoff, Gahyun Kim, Jan D. Lanzer, Matthias Kretzler, Ricardo O. Ramirez Flores, Julio Saez-Rodriguez

## Abstract

Fibrotic scarring is a common response to tissue injury. Repeated or severe insults can cause fibrosis, leading to excessive extracellular matrix deposition and a substantial clinical risk of organ dysfunction. Despite its high prevalence, few therapeutic options exist, and fibrotic diseases collectively represent a major global health burden. Fibrotic diseases affect virtually all organs, yet they have been explored mainly in isolation for each organ. As a result, proposed shared fibrotic mechanisms are often based on indirect comparisons between independent datasets rather than on a unified, systematic, cross-organ meta-analysis. To overcome this gap, we conducted a large-scale meta-analysis of single-cell transcriptomic data from healthy and fibrotic human tissues to identify both shared and organ-specific transcriptomic profiles. We constructed a single-cell fibrosis atlas of over five million cells from 20 studies, covering more than 25 disease etiologies affecting the heart, liver, kidney, and lung. Through systematic comparison of these datasets, we identified organ-specific as well as cross-organ fibrosis-associated gene expression profiles in major cell types and defined disease fibroblast subpopulations with excessive extracellular matrix production. These analyses revealed a conserved fibrotic response shared across tissues. Our analysis spans global comparisons of fibrosis-associated changes in cellular composition and predictive disease signatures to detailed examinations of individual genes, transcription factors, and intercellular communication patterns observed in fibrotic diseases across organs. We provide our cross-organ integration as a user-friendly open resource for investigating fibrotic diseases across organs. This resource will enable an accelerated discovery of disease mechanisms and faster development of broadly effective antifibrotic strategies in the future.

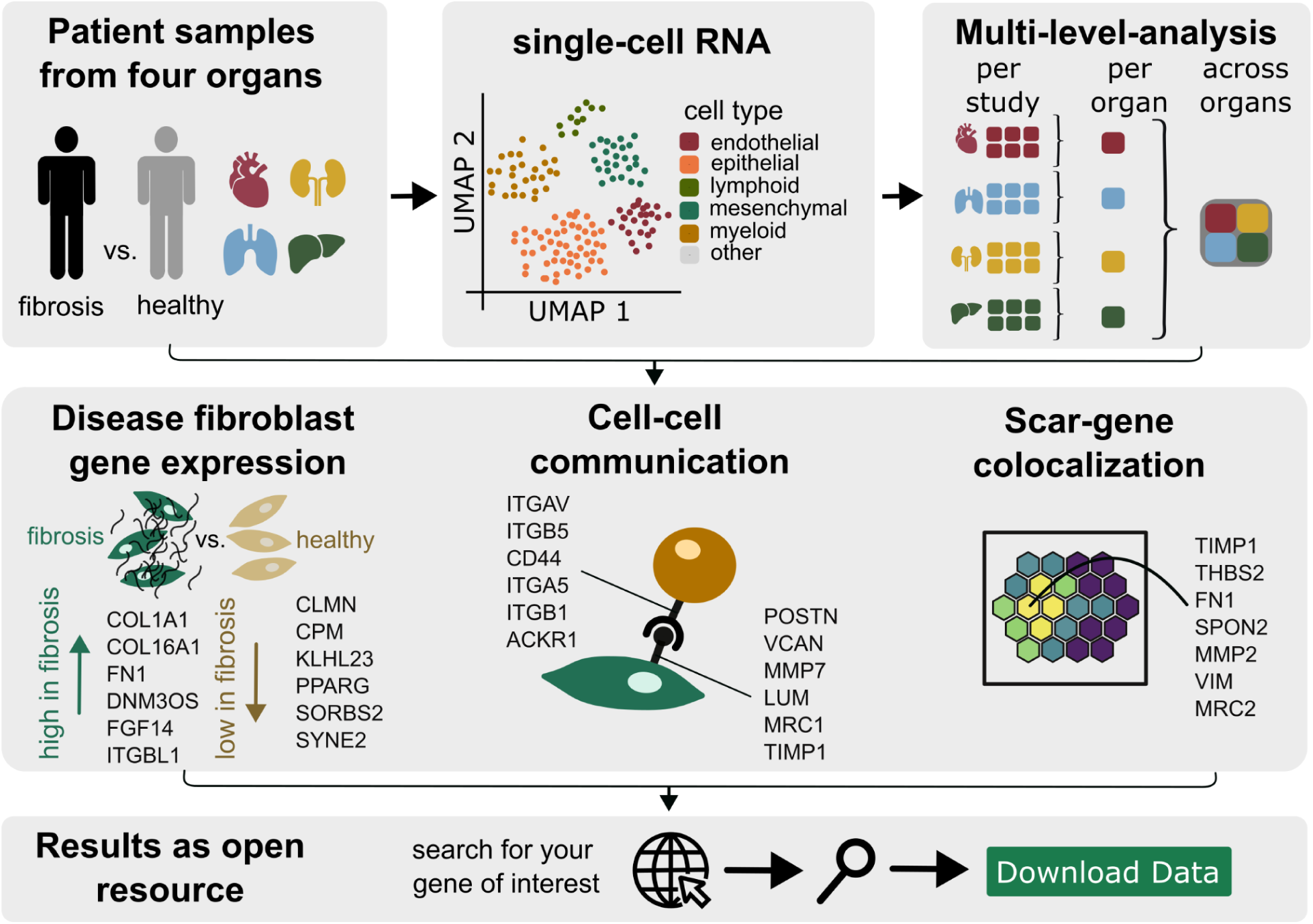
Graphical abstract.

## Introduction

Fibrosis is characterized by progressive, irreversible extracellular matrix (ECM) deposition^1^. Ultimately, excessive fibrotic tissue can lead to tissue architecture disruption, organ dysfunction, and, in the worst case, organ failure. Fibrotic diseases can affect almost any tissue, and they are estimated to be responsible for 17.8% – 35.4% of global deaths^2^. The initial physiological role of fibrotic tissue formation is to support wound healing. However, repeated or severe tissue injuries cause excessive fibrotic tissue production, resulting in a deterioration of organ structure and function^3^. These injuries can be caused by infections, exposure to environmental irritants, repeated mechanical trauma, genetic predispositions, or autoimmune diseases, among others^4^.

Even though fibrotic diseases are diverse in cause and location in the human body, they share common mechanisms^3–6^. These include the activation of myofibroblasts, activated fibroblasts with contractile abilities that deposit large amounts of ECM proteins. While their existence may be shared across organs, several cellular origins of myofibroblasts have been reported, including pericytes, fibroblasts, epithelial cells, endothelial cells, and adipocytes^4,7^, and it is unclear to what extent they share a common transcriptomic profile. Other common mechanisms across organs include *TGF-β* signaling, which has been identified as a major pro-fibrotic modulator of tissue scarring in the liver, lung, kidney, skin, and heart. Consequently, several studies and clinical trials have been investigating the pathway as a potential drug target in organs including the heart, lung, kidney, and liver^5,8,9^. Given these and other shared features, it is reasonable to hypothesize that antifibrotic therapies targeting common mechanisms could be effective across organs. However, only three anti-fibrotic drugs, pirfenidone, nintedanib, and nerandomilast, have been approved to date as treatment options for pulmonary fibrosis^6,10^, and no anti-fibrotic drug is approved for more than one tissue^6^.

Most biological findings about fibrotic diseases stem from single studies that investigate a specific disease etiology in a single, isolated tissue at a time (Table 1). These studies mainly focus on cataloguing different cell states, yielding highly specific cell groups that are difficult to compare across datasets, as they are often defined by only one or two marker genes, lack robustness, and offer limited meaningful comparisons^11^. While few studies have directly compared fibroblasts across tissues^12–14^, they have not accounted for other cell types that also play a role in fibrotic diseases. Without a direct comparison of patient data from multiple studies, tissues, and diseases, it remains unclear to what extent these single findings are transferable between fibrotic diseases, both within and across tissues.

**Table 1.**
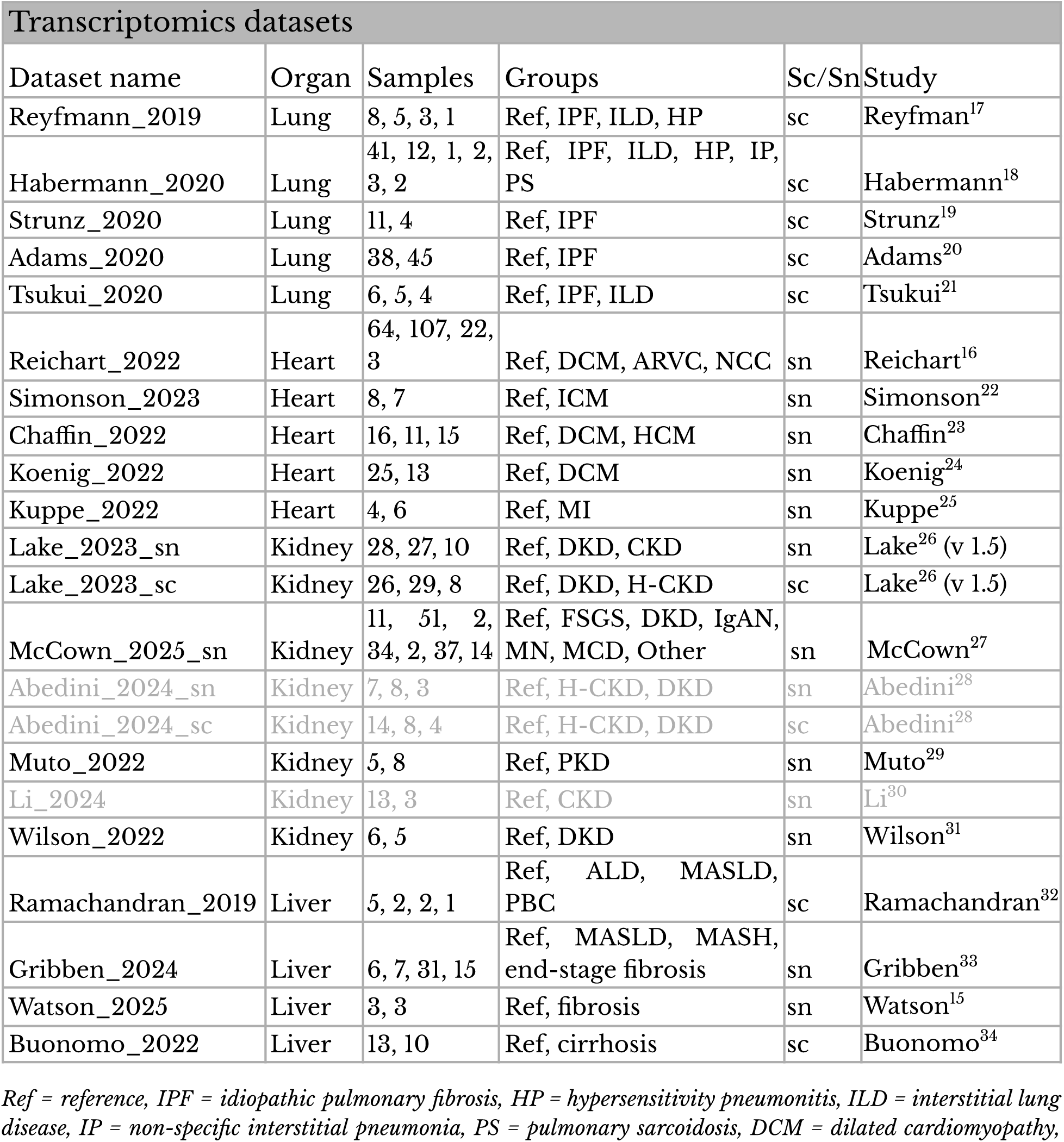
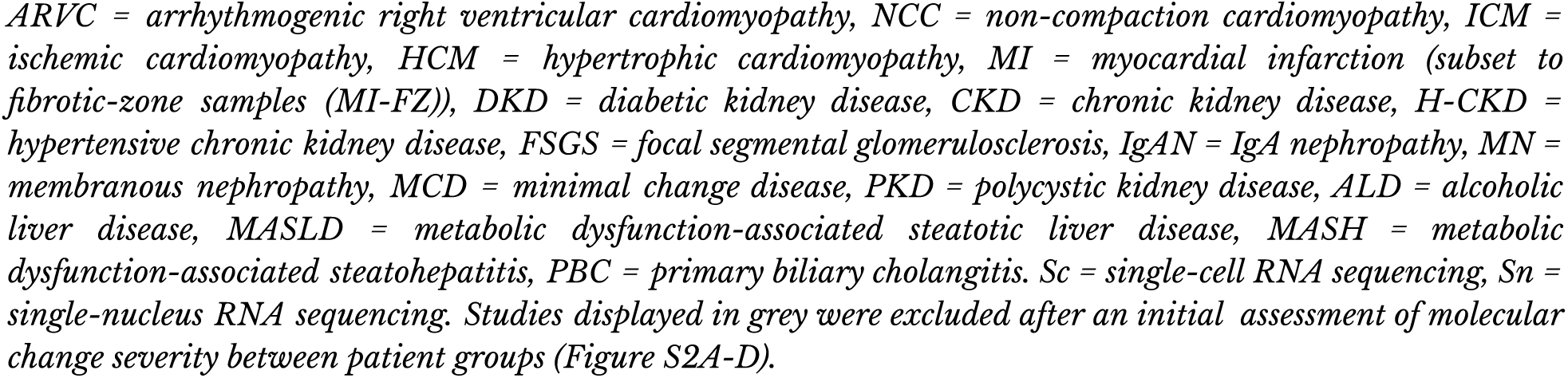
Overview of curated single-cell and single-nucleus RNA datasets.

As the tissues most affected by fibrotic diseases^2^, we compare the molecular phenotype of diverse fibrotic diseases in human heart, liver, lung, and kidney samples. We perform a large-scale meta-analysis of single-cell transcriptomic datasets (Table 1) to extract shared and unique mechanisms across five broad cell type lineages that are present in most organs. We assess how strongly a shared fibrotic gene program emerges across studies from the same organ and, in the next step, across different organs. Our analysis ranges from global comparisons of cellular composition and predictive disease signatures to detailed insights into individual genes, transcription factors, and intercellular communication events observed in fibrotic diseases across organs. Alongside this manuscript, we publish a user-friendly resource (https://organfibrosis.saezlab.org/) that makes all results easily accessible and searchable to answer specific questions.

## Results

### Data curation for a cross-organ comparison of fibrotic diseases

We queried public databases to select several single-nucleus and single-cell RNA (snRNA / scRNA) sequencing datasets profiling diverse fibrotic processes (Table 1). We focused on four tissues majorly affected by fibrotic diseases^2^ – heart, lung, kidney, and liver (Figure 1A) and restricted our search to studies with publicly available human fibrotic disease samples, regardless of their fibrotic disease etiology, and healthy reference samples. Studies with both snRNA and scRNA data were split into two distinct datasets.

**Figure 1.**
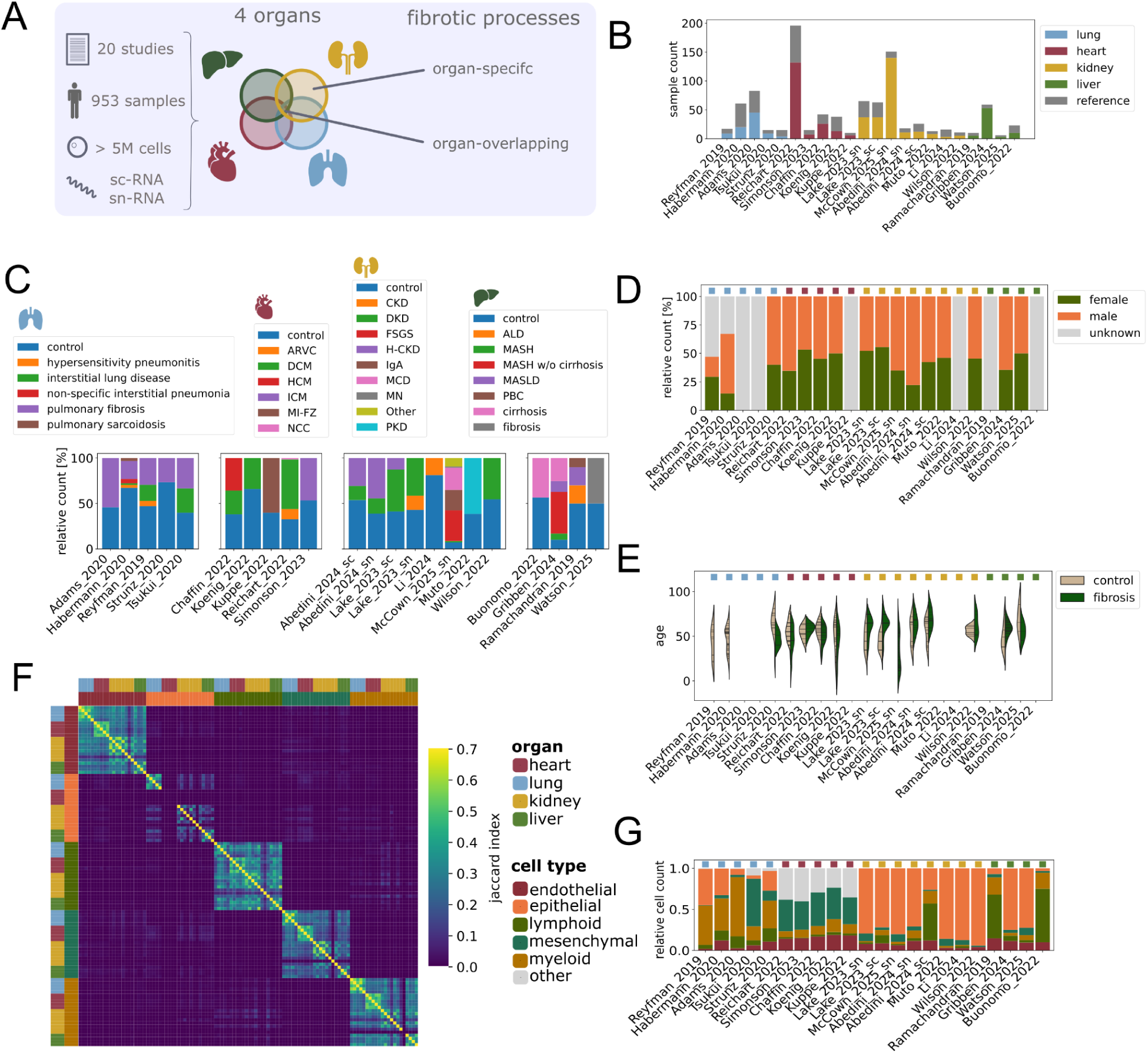
Overview of curated datasets. (A) Study overview – this study combines 20 human single-cell and single-nucleus RNA sequencing datasets from the heart, lung, liver, and kidney. Organ-specific and overlapping fibrotic processes are identified. (B) Number of fibrosis (color) or reference (grey) samples per dataset. Studies are colored by their tissue of origin. (C) Relative number of samples per disease etiologies for each study (from left to right: lung, heart, kidney, liver). (D) Gender distribution per dataset. Colored boxes above indicate the tissue of origin. (E) Age distribution per dataset. Colored boxes above indicate the tissue of origin. (F) Jaccard Index of cell type marker gene (FDR < 0.01 and logFC > 2) overlap. Each row and column reflects one study. Outer colors indicate which organ the study comes from; inner colors indicate the cell type. (G) Relative cell type distribution per dataset. Colored boxes above indicate the tissue of origin.

Overall, 20 scRNA or snRNA studies were curated (heart: 5, lung: 5, kidney: 6, liver: 4). The study size varied from small-scale datasets with three disease and three reference samples (Watson_2025^15^, liver) to large-scale studies with 132 disease and 64 reference samples (Reichart_2022^16^, heart) (Figure 1B). Furthermore, the number of disease etiologies covered per study varied (Figure 1C). In lung datasets, idiopathic pulmonary fibrosis (IPF) was the best-described disease, with 71 samples (vs. 16 from other disease etiologies). At the same time, dilated cardiomyopathy (DCM) was the most common disease in the heart, with 131 samples (vs. 53 from other disease etiologies). In the other organs, disease etiologies were more spread, with the most predominant disease in the kidney being diabetic kidney disease (DKD, 70 vs. 183 samples from other disease etiologies) and in the liver metabolic dysfunction-associated steatohepatitis (MASH, 31 vs. 44 samples from other etiologies). We collected a total of 953 samples (heart: 301, lung: 191, kidney: 363, liver: 98) and 5,109,160 cells (heart: 2,366,669; lung: 684,316; kidney: 1,832,416; liver: 225,759). We also collected metadata wherever available and harmonized it. In studies with available gender information (16/22 datasets), we found a slight gender bias toward the male sex (avg. 43 ± 9% female; Figure 1D). The average age of patients was highest in the liver (fibrosis: 58 yrs., reference: 56 yrs.), followed by the heart (fibrosis: 54 yrs., reference: 47 yrs.), lung (fibrosis: 47 yrs., reference: 49 yrs.), and finally the kidney (fibrosis: 45 yrs., reference: 50 yrs.) (Figure 1E). Seven out of the twenty studies did not include age information at all, and three studies only included it for the control or fibrosis population. In conclusion, our curation efforts resulted in a dataset that captures a diverse range of fibrotic disease etiologies.

In addition to the reported biological diversity, we also found technical variability across studies and organs. Some studies had lower total RNA and feature counts per cell than others (Figure S1A, B), including Strunz_2020 in the lung, McCown_2025_sn, Abedini_2024_sc, Muto_2022, and Wilson_2022 in the kidney, and Buonomo_2022 in the liver. Overall, the kidney was the organ with the lowest RNA counts (average counts per cell kidney: 2624, heart: 3323, liver: 4986, lung: 5884) and feature counts (average feature counts per cell kidney: 884, heart: 1494, liver: 1968, lung: 1846). On the other hand, a low average cell number per sample was found in the liver, and an especially high cell number per sample in the heart (liver: 9035, lung: 20,700, kidney: 35,515, heart: 56,866). As expected, mitochondrial gene counts were consistently higher in scRNA than snRNA studies (Figure S1C) and were especially high in the kidney.

To enable a comparison across tissues, we grouped cells into five broad categories based on the author’s annotations: endothelial-, epithelial-, mesenchymal-, lymphoid-, and myeloid cells (Suppl. Table 1). In datasets without such classifications, we applied a cell type label transfer (see methods), which was only required for two kidney studies (Muto_2022^29^, Wilson_2022^31^). Cell types without clear equivalents across tissues (e.g., cardiomyocytes) were labeled as ‘other’ and were excluded from this analysis. Although many cell types are shared across organs, cells of the same type can fulfill distinct functions depending on tissue context. To assess how comparable cell identity and characteristics are across organs, and to flag potential misannotations, we quantified their degree of similarity by comparing their marker gene expression profiles (Figure 1F). The same cell types across organs showed higher marker gene similarity than different cell types within the same organ (within cell type avg. Jaccard index: 0.22; between cell types: 0.01). Moreover, cells from the same cell type and organ showed, in turn, greater similarity (avg. Jaccard index: 0.34) than across organs (avg. Jaccard index: 0.20), suggesting that within a cell lineage, organ-specific differences exist. Despite similarities in gene expression, cell type abundances varied substantially across organs (Figure 1G). While myeloid cells accounted for an average of 42.0% of lung samples, they were sparsely represented in kidney samples (avg. 2.6%). Furthermore, heart samples lacked epithelial cells, whereas they made up, on average, 82.2% of all kidney cells. In addition, some studies experimentally enriched for cell types of interest, which impacted the full distribution of cell types. Consequently, the representation of each cell type group varied from organ to organ (Figure S1D, E), with the heart having the most samples with more than 50 cell types per group. Together, these observations indicate that major cellular lineages maintain comparable molecular profiles across organs, even though they are captured at different proportions.

### Fibrotic disease programs within tissues

First, we aimed to extract robust fibrotic disease signals within organs. To ensure that each included dataset displays a strong molecular difference between healthy and fibrotic patients, we evaluated the magnitude of transcriptomic change from reference to fibrotic samples for each study and cell type. To do so, we applied ScDist^35^, a method that estimates the distance between cell populations and that can be used to measure which cell types contribute most to the change between two conditions. Within each study, we quantified molecular changes per cell type between healthy and fibrosis patient groups (Suppl. Figure 2A-D). As we focused on common fibrotic processes across disease etiologies, we grouped all fibrosis patients and contrasted them with reference patients in their study. Three kidney studies displayed low to no change between patient groups and were therefore excluded from further analysis (Figure S2A-D, excluded datasets: Abedini_2024_sc^28^, Abedini_2024_sn^28^, and Li_2024^30^). To test if these results are confounded by study size, we confirmed that there is no clear relationship between cell counts per study and estimated distances (Figure S2E).

We next investigated cell type proportion changes for each study and tested if similar patterns were observed within each organ. Compositional analysis with linear mixed models (see methods) revealed distinct changes in proportion across tissues (Figure 2A). In the lung, we observed an increase in mesenchymal cells in fibrotic samples (adj. pval = 0.015), while in the heart and kidney, we observed an increase in lymphoid cells (heart adj. pval = 0.001; kidney adj. pval = 0.002). While heart studies showed the highest agreement between studies, we overall detected a high study-to-study variability in cell type proportion changes, even at this broad cell type level (average percent study variance heart: 8.7%, lung: 27.5%, kidney: 33.8%, liver: 32.9%). These results suggested that cell proportions differed not only between patients and organs, but also that there was no uniform abundance change in disease.

**Figure 2.**
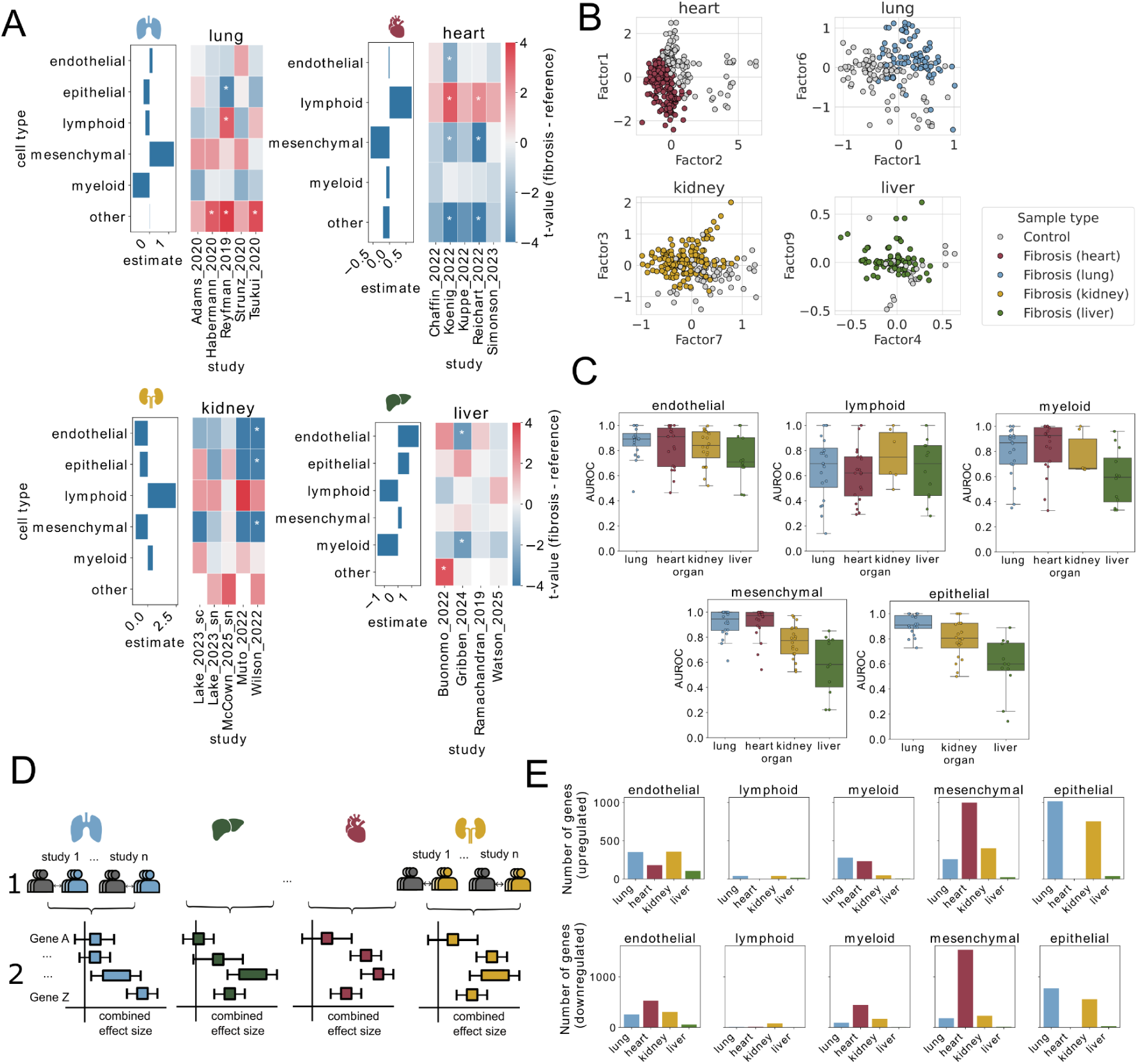
Fibrotic processes within organs. (A) Differential compositional analysis: T-statistics (heatmap) and estimate (bars) of the difference between fibrotic and non-fibrotic samples using t-tests and linear mixed models, respectively. Stars denote an adj. p-value < 0.05. (B) Latent factors from multicellular factor models that most strongly associated with patients’ disease status per organ model. Each dot is one sample; grey samples are reference samples, colored samples are fibrosis samples. (C) Disease score predictions within tissues per cell type and organ. Each dot represents one prediction score from one study to another. (D) Method overview – Differential gene expression of fibrosis vs. reference samples is assessed per study (level 1) and is then summarized into an organ effect size (level 2), resulting in a combined effect size per gene and organ. (E) Number of upregulated (upper panel) or downregulated (lower panel) organ consensus genes for each cell type. A gene was defined as an upregulated organ consensus gene with a combined effect size > 0.5 and a lower confidence interval > 0 or as downregulated with a combined effect size < –0.5 and an upper confidence interval of < 0.

Next, we performed an unsupervised analysis of all samples and studies within an organ to identify a shared signal of fibrosis and evaluate the degree to which each organ displays common fibrotic molecular signature across studies. Here, we employed a multicellular factor analysis model^36^, a method that can extract common axes of variation of patient samples across cell types. We reasoned that if a factor consistently distinguishes fibrotic from healthy samples across multiple studies, this factor reflects a shared molecular signature of fibrosis within an organ. For heart, kidney, and lung, at least one factor captured a molecular program that could clearly separate fibrosis from reference patients in at least four out of five studies, which hints at a common fibrotic process within these organs (Figure 2B, Figure S3A). In contrast, no factor captured a representative program for all liver studies, suggesting either strong biological or technical differences between the studies.

Given that there were common disease-associated multicellular programs in three organs, we investigated which cell types contributed to this convergence. To specifically quantify the extent of shared gene regulation across studies within each cell type, we employed a disease-score prediction model (see methods). Briefly, the disease score prediction uses the top differentially expressed genes from one cell type in one study to predict the disease status in patients from an independent study. High predictive performance suggests similar fibrotic gene regulation between these two studies. Overall, heart studies displayed the highest within-organ prediction scores (median AUROC: 0.92), followed by lung (median AUROC: 0.89), and kidney (median AUROC: 0.77) (Figure 2C). Consistent with results from the multicellular factor analysis in the liver, comparisons between liver studies revealed the lowest degree of shared transcriptional signal (median AUROC: 0.60). We also observed differences between cell types: The highest within-organ prediction performance was observed for mesenchymal cell populations in heart and lung (median AUROC: heart = 0.97, lung = 0.94). Additionally, lymphoid cells consistently exhibited minimal overlap in signal across studies and cell type comparisons (median AUROC heart = 0.62, lung = 0.70, kidney = 0.75, liver = 0.70), whereas endothelial cells displayed the highest median overlap (median AUROC heart = 0.91, lung = 0.89, kidney = 0.84, liver = 0.70). Overall, these analyses demonstrate that, despite substantial variation in cell type composition across studies, a common fibrosis signature can be detected in the lung, heart, and kidney.

### Identification of shared gene programs per tissue

Having established that shared transcriptional signals exist within individual organs, we next performed a meta-analysis of differentially expressed genes to identify those contributing to the common molecular profile of fibrotic tissues. We first extracted differentially expressed genes between reference and fibrosis patients per study. Using a random-effects model with inverse-variance weighting, these results were then combined into a unified organ effect size that reflects the consistency of gene regulation within individual organs. Genes with significant organ effect sizes will from hereon be referred to as organ-consensus genes; genes regulated similarly across organs with significant cross-organ effect sizes will be termed cross-organ consensus genes (Figure 2D).

This methodology enabled a ranking of the most up– and downregulated genes per organ. Highest confidence upregulated genes (present in at least three studies, ranked by organ effect size) in mesenchymal cells, for instance, consisted of *SEZ6L, FAM155A, FOSB* in the heart; *POSTN, MXRA5, THY1* in the lung; *IGKC, C3, PIGR* in the kidney; and *MOXD1, CCDC80, LXN* in the liver. In endothelial cells, another cell type that displayed high agreement within organs, the top upregulated genes were *AC093817, NPPB,* and *NPPA* in the heart; *APLNR, SPP1,* and *COL15A1* in the lung, *VCAN, SYT1,* and *MSC-AS1* in the kidney; and *FABP4, ZNF385D,* and *VWF* in the liver. A full list of the top ten upregulated genes per organ and cell type can be found in Suppl. Table 2. Counting the number of organ-consensus genes based on their organ effect sizes and confidence intervals (see methods) revealed different numbers of commonly regulated genes per cell type and organ (Figure 2E): While there were on average only 36.8 organ-consensus genes per cell type in liver studies, other organs shared more upregulated genes (mean in heart: 353.8, lung: 388.8, kidney: 319.8). Similarly, the average number of downregulated organ-consensus genes was much lower in the liver (19.4) than in the other organs (heart: 630.25, lung: 267.4, kidney: 262.6). With an exception to the liver, the number of organ-consensus genes generally aligned with the number of differentially expressed genes detected between fibrosis and reference samples per cell type (Figure S4A): For instance, heart studies showed the most number of differentially expressed genes in mesenchymal cells (avg. DEGs in mes. cells heart: 3008, Figure S4A) and similarly exhibited the most amount of organ-consensus genes in mesenchymal cells (upregulated: 1000, downregulated: 1536) (Figure 2E). In contrast, liver studies yielded very few organ-consensus genes, indicating limited agreement in gene regulation across datasets despite the presence of differentially expressed genes within individual studies. Overall, these results suggest that there is a part of the fibrotic program that appears to be shared within most tissues, primarily found in endothelial, mesenchymal, and epithelial cells.

### Tissue comparison

Thus far, our analysis has focused on tissue-specific regulation of fibrotic genes. Similar to within-tissue analyses, we also quantified the degree of conservation across tissues. We first compared the overlap of organ-consensus genes between organs for each cell type. This analysis revealed that organ-consensus genes were largely organ-restricted, with an average of 87.2% of significantly regulated genes being organ-specific. (Figure 3A). While the differences between cell types were not strong, most organ-consensus genes shared between organs were found within endothelial cells (unique genes up: 79.9%, down 83.3%) and mesenchymal cells (unique genes up: 77.1%, down 85.2%), whereas lymphoid cells displayed most genes specific to one organ (up: 92.9%, down: 100%). Of note, no single organ-consensus gene in any of the five cell type groups was up– or down-regulated in all four organs (Figure S4B & C).

**Figure 3.**
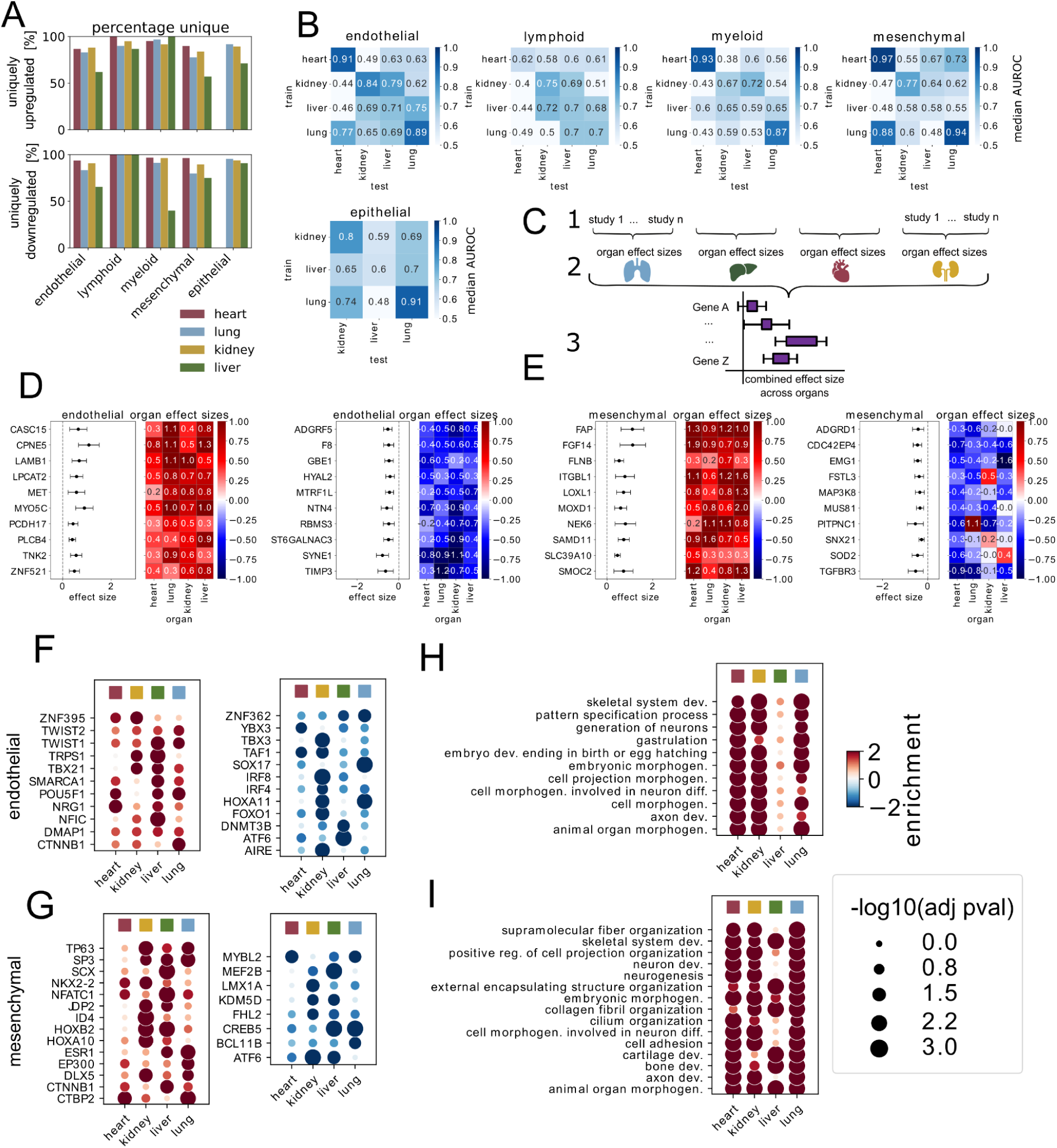
Fibrotic processes across organs. (A) Percentage of upregulated (top) or downregulated (bottom) organ consensus genes that are unique to the organ in which they were identified. (B) Median AUROC values for disease score predictions across tissues and cell types. (C) Organ effect sizes (shown in Figure 2D) were summarized to retrieve a cross-organ effect size per gene. (D) Top ten up– (left) and down-regulated (right) genes across tissues in endothelial cells based on their cross-organ effect sizes. Boxplot on the left displays the cross-organ effect size, whereas heatmap on the right displays the organ-effect sizes for each organ. (E) Top ten up– (left) and down-regulated (right) genes across tissues in mesenchymal cells based on their cross-organ effect sizes. Boxplot on the left displays the cross-organ effect size, whereas heatmap on the right displays the organ-effect sizes for each organ. (F) Most commonly up– (left) and down-regulated (right) transcription factor activities in fibrosis across organs in endothelial cells and (G) mesenchymal cells. (H) Most commonly up-regulated go-terms in fibrosis across organs in endothelial cells and (I) mesenchymal cells. For F-I: Dots show the enrichment score (positive: upregulated in fibrosis), while sizes show the –log10 of the adjusted p-values of ULM enrichments.

Because direct comparisons of differentially expressed genes can be overly conservative as they rely on specific cutoffs^37^, we evaluated whether organs nonetheless exhibit a similar global transcriptional shift by applying our disease prediction models across organs (Figure 3B): Here, mesenchymal cells displayed good prediction values between heart and lung, suggesting similar transcriptional regulation in fibrosis between these two organs. On the other hand, the organ comparison of myeloid cells revealed a low degree of shared transcriptional signal, even though each tissue displayed coherent myeloid gene programs on its own. This pattern may indicate differences in the regulation of myeloid cells in fibrosis across tissues and is consistent with our observation that immune-related programs generally exhibit low cross-organ overlap, as also reflected in the poor predictive performance observed for lymphoid cells. Furthermore, we detected similarities in endothelial cell programs between the kidney and liver, and heart and lung, as well as epithelial similarities between the kidney and lung. Interestingly, the model maintained similar performance when organ-consensus genes were excluded from the model (Figure S5). This observation indicates that fibrotic gene regulation is not solely driven by genes that pass a significance threshold. Instead, despite differences in which genes are individually significant, the broader pattern is consistent, with up– and downregulated genes in one organ tending to be up– and downregulated in others, respectively. These analyses revealed that different organs and cell types exhibit varying degrees of shared fibrotic molecular signatures, with the highest directionality concordance observed in mesenchymal, endothelial, and epithelial cells. Furthermore, the heart and lungs, and to a lesser extent the kidney, showed greater agreement between studies than the liver.

### Common aspects of fibrosis across tissues in endothelial and mesenchymal cells

Although most of the strongly up– or downregulated genes within individual tissues appeared to be organ-specific, our predictive models indicated that many genes are nevertheless regulated in the same direction across tissues. To identify which genes show the highest agreement in fibrotic regulation, we applied a second random-effects model to extract genes that are similarly regulated across organs. In this model, we combined the organ-effect sizes into a single cross-organ summary effect size (Figure 3C). We focused on endothelial and mesenchymal cells because prediction models indicated that these cell types showed the greatest degree of shared gene regulation across organs. In endothelial cells, some of the genes with the ten highest cross-organ effect sizes were genes that have already been reported to be upregulated in the context of fibrotic diseases (Figure 3D), including Laminin subunit beta-1 (*LAMB1*)^38,39^, Protocadherin-17 (*PCDH17*)^40^, or Zinc finger protein 521 (*ZNF521)*^41^. Similarly, the top ten upregulated genes in mesenchymal cells contained many established pro-fibrotic genes (Figure 3E). For instance, SPARC-related modular calcium-binding protein 2 (*SMOC2*), a protein that promotes matrix assembly, has been implicated in fibrotic diseases and shown to be a potential therapeutic target across all four organs investigated in this study^42–45^. Likewise, Lysyl oxidase homolog 1 (*LOXL1*), a protein that mediates the cross-linking of collagens and elastins, has been reported to contribute to matrix stiffening and fibrosis in experimental models of lung^46^ and liver^47^ fibrosis. Similarly, DBH-like monooxygenase protein 1 (*MOXD1*) might be worth further exploration, as it has been therapeutically investigated in the liver^48^ and kidney^49^, but not in other organs to our knowledge. Other genes with high cross-organ effect sizes have been less well described, such as fibroblast growth factor 14 (*FGF14*).

From organ effect sizes, we also inferred transcription factor (TF) activities per organ using CollectTRI^50^, a curated collection of TF-targets, and identified the most commonly upregulated TFs based on the up– or downregulation of the genes they regulate across organs. In endothelial cells, well-explored profibrotic TFs such as *TWIST1*^51^, *TWIST2*^52^, or *CTNNB1*^53^ (β-Catenin) were identified as common profibrotic factors, as well as the anti-fibrotic factor *FOXO1*^54^. On the other hand, several TFs with no specified role in fibrogenesis were also revealed, including *SMARCA1, DMAP1, or ZNF362* (Figure 3F). Similarly, commonly upregulated mesenchymal TFs included known profibrotic factors (including *SP3*^55^, *SCX*^56^, *NFATC1*^57^, *EP300*^58^) and several TFs that have not been implicated in fibrotic diseases (Figure 3G). Several transcription factors associated with developmental gene programs, such as several homeobox genes (*HOXB2, HOXA10*), the stem cell factor *POU5F*, or chromatin remodeling *SMARCA1,* emerged in the analysis. This association with development was further supported by the functional characterization of upregulated genes in each organ. Both mesenchymal and endothelial cells showed reactivation of developmental and morphogenetic programs. Endothelial cells displayed an enrichment of embryonic morphogenesis, gastrulation, pattern specification, and neuronal morphogenesis pathways, reflecting developmental reprogramming and vascular remodeling (Figure 3H). Mesenchymal cells showed strong upregulation of pathways involved in supramolecular fiber and collagen fibril organization, skeletal and cartilage development, and cell adhesion, consistent with enhanced extracellular matrix production and structural remodeling. In addition, enrichment of neuronal development and axonogenesis pathways points to activation of projection-related programs (Figure 3I). Together, these findings suggest that both endothelial and mesenchymal cells undergo coordinated rewiring in association with fibrosis, with developmental and remodeling processes activated.

### Disease fibroblasts converge in a common fibrotic gene profile

Given the strong similarities between mesenchymal cells across organs, we examined this population further. Several studies have previously reported specific mesenchymal cell states to play an important role in fibrotic diseases, such as *CTHRC1*+ fibroblasts^21^ or *SSTR2*+ pericytes^59^ in the lung, *FAP/POSTN*+ fibroblasts in the heart^60^, *NKD2*+ fibroblasts in the kidney^7^, *CXCL14*+ cells in the intestine^61^, or *PDGFRα*+ collagen-producing myofibroblasts in the liver^32^. Examining the studies separately makes it difficult to assess how these cell states compare. However, their central involvement in fibrotic disease underscores the importance of understanding how they relate to each other. To identify disease-relevant cell states in our datasets, we integrated the mesenchymal cell population per organ and identified a disease-associated cluster by compositional analysis (Figure 4A, Figure S6A-C). Similar to our analysis across cell types, we extracted organ-consensus and cross-organ-consensus genes from disease fibroblast state markers. For this analysis, disease fibroblasts were contrasted with the remaining mesenchymal cell populations in each study. These results were then summarized into organ– and cross-organ effect sizes. Overall, we observed the assimilation of gene programs of disease fibroblasts across organs: organ-consensus genes were less organ-specific (Figure 4B), leading to more organ-overlapping genes (Figure 4C). The top ten cross-organ consensus genes were mostly genes that are already known to play a role in fibrosis, and partly genes that were not yet investigated as thoroughly (Figure 4D). For instance, disease-associated fibroblasts displayed many collagens among their most commonly upregulated genes. On the other hand, one of the most down-regulated genes in disease fibroblasts across organs was Peroxisome proliferator-activated receptor gamma (*PPARG*), a gene known for its anti-fibrotic effects^5^. Agonists are therefore currently investigated for the treatment of fibrotic disease in many organs^62–65^.

**Figure 4.**
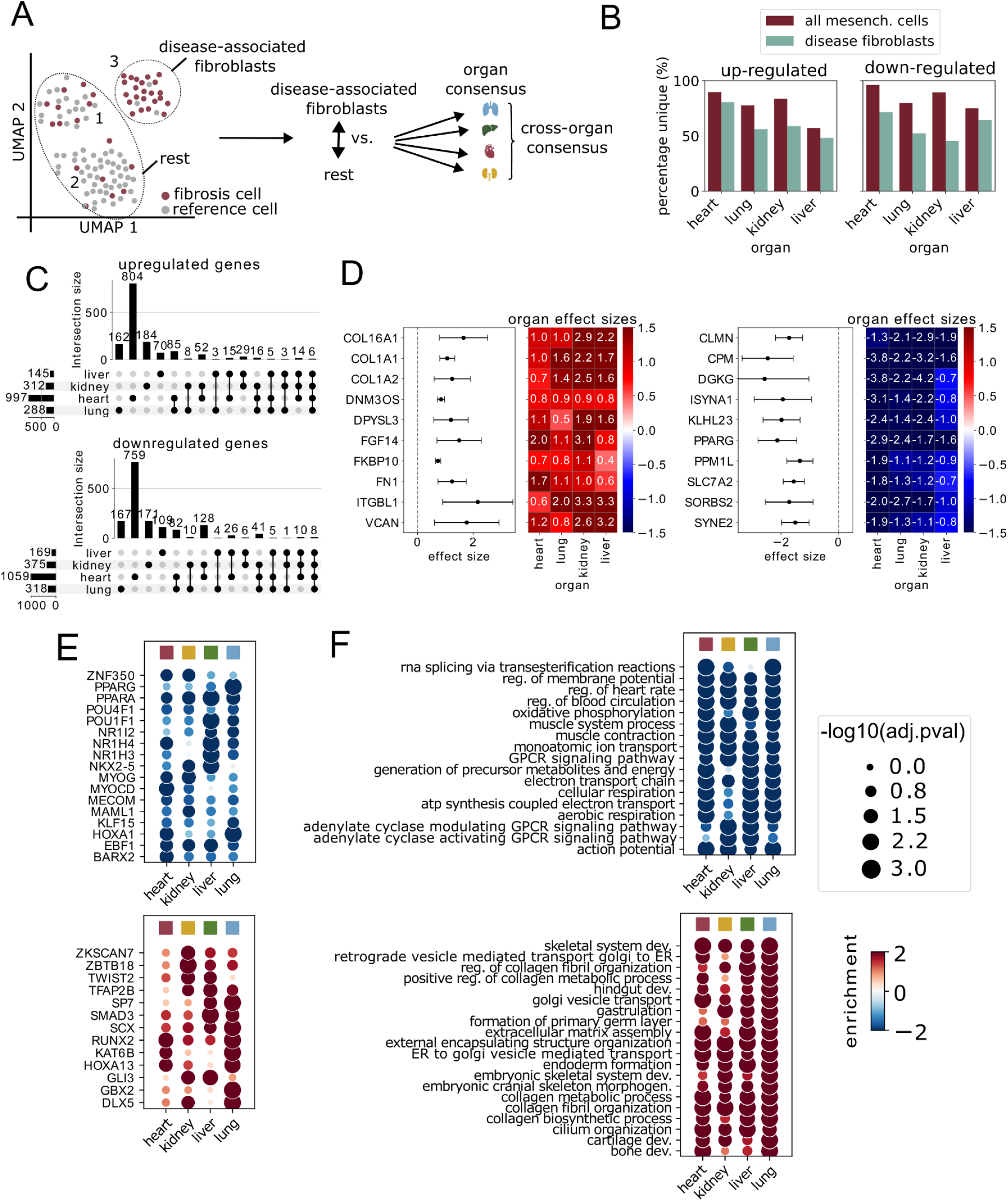
Disease-associated fibroblasts. (A) Schematic of methodology of disease-associated fibroblast identification: Mesenchymal cells were integrated into one latent space per organ and subsequently clustered (here: cluster 1 – 3). Through a compositional analysis of disease vs. reference cells, disease-associated clusters were identified (here: cluster 3). Disease-associated fibroblasts were contrasted with other mesenchymal cells (“rest”) per study and were again summarized into organ– and cross-organ-level effect sizes, as displayed in Figure 3D. (B) Percentage of upregulated (left) or downregulated (right) organ-consensus genes in mesenchymal cells (red) vs. disease fibroblasts (mint) that are unique to the organ they were identified in. (C) Absolute number of up-regulated (upper panel) and down-regulated (lower panel) disease-associated fibroblast organ consensus genes and their overlap between organs. (D) Top ten up– (left) and down-regulated (right) genes across tissues in mesenchymal cells based on their cross-organ effect sizes. Boxplot on the left displays the cross-organ effect size, whereas heatmap on the right displays the organ-effect sizes for each organ. (D) Top predicted TFs to be active (upper panel) or inactive (lower panel) across tissues in disease-associated fibroblasts. (E) Common go-terms across tissues in disease-associated fibroblasts. For E-F: Dots show the enrichment score (positive: upregulated in disease-associated fibroblasts), while sizes show the –log10 of the adjusted p-values of ULM enrichments.

We also investigated TFs predicted to be active (Figure 4E). The list of downregulated TFs included many factors that have shown anti-fibrotic effects. For instance, the introduction of an agonist or upregulation of *PPARG*^66^, *PPARA*^67^, *NR1I2*^68^, *NR1H4*^69^, and *NR1H3*^70^ has all been shown to have anti-fibrotic effects in several experimental models. Upregulated TFs included well-known pro-fibrotic TFs such as *SMAD3*^71^ and *SCX*^72^, as well as many that are not known to play a role in fibrosis, including *ZKSCAN7*, *TFA2B*, and *HOXA13*. Gene Ontology enrichment analysis revealed upregulation of developmental, extracellular matrix, and secretory pathways (Figure 4F). In particular, pathways associated with collagen metabolism, fibril organization, extracellular matrix assembly, and skeletal-, bone-, and cartilage development were prominently enriched, indicating enhanced matrix synthesis and tissue remodeling consistent with fibrotic activation. Increased vesicle-mediated transport between the Golgi apparatus and endoplasmic reticulum further implied heightened secretory and protein-processing activity. In contrast, downregulated GO categories included oxidative phosphorylation, cellular respiration, and ion transport, indicating an alteration in energy metabolism and a shift towards structural and biosynthetic programs characteristic of fibrotic remodeling. Overall, we observed a convergence of fibrotic signature genes across organs when specifically investigating the changes in disease-enriched fibroblasts, reflected by high similarities in regulated genes, transcription factors, and biological functions across tissues.

### Cell-cell communication similarities between organs

While fibroblasts are often said to cause the pathogenic phenotype in fibrotic patients, they are most likely acting together with other cell types in their surroundings. Therefore, we analyzed which cell-cell communication events could lead to or sustain the fibrotic state of the tissues. After combining differentially expressed genes per organ with a mixed effects model, we identified ligand-receptor pairs that were commonly regulated in fibrosis across studies within one organ. We found on average slightly more up– (heart: 52.7, lung: 118.1, liver: 28.6, kidney: 155.8) than down-regulated (heart: 55.6, lung: 33.6, liver: 33.2, kidney: 33.0) communication events in fibrosis compared to reference samples (Figure S7A & B). In the heart, the largest fraction of communication events was found between mesenchymal and endothelial cells (26.5%) and within the mesenchymal cell population (26.5%). In the other three organs, the epithelial cell population additionally made up a large fraction of the communication events, especially in the kidney and liver, in combination with endothelial (kidney: 19.7%, liver: 11.2%, lung: 9.2%) and mesenchymal cells (kidney: 17.1%, liver: 14.8%, lung 12.8%) (Figure 5A). When filtering specifically for ligand-receptor pairs that were upregulated in fibrotic samples across all four organs, we retrieved nine ligand-receptor pairs (Figure 5B). These mostly entailed well-known upregulated genes in fibrotic conditions, such as collagen 1A1 (*COL1A1*)^73^, fibronectin (*FN1*)^74,75^, periostin (*POSTN*)^76–79^, prolyl endopeptidase FAP (*FAP*)^80,81^, or versican (*VCAN*)^82,83^.

**Figure 5.**
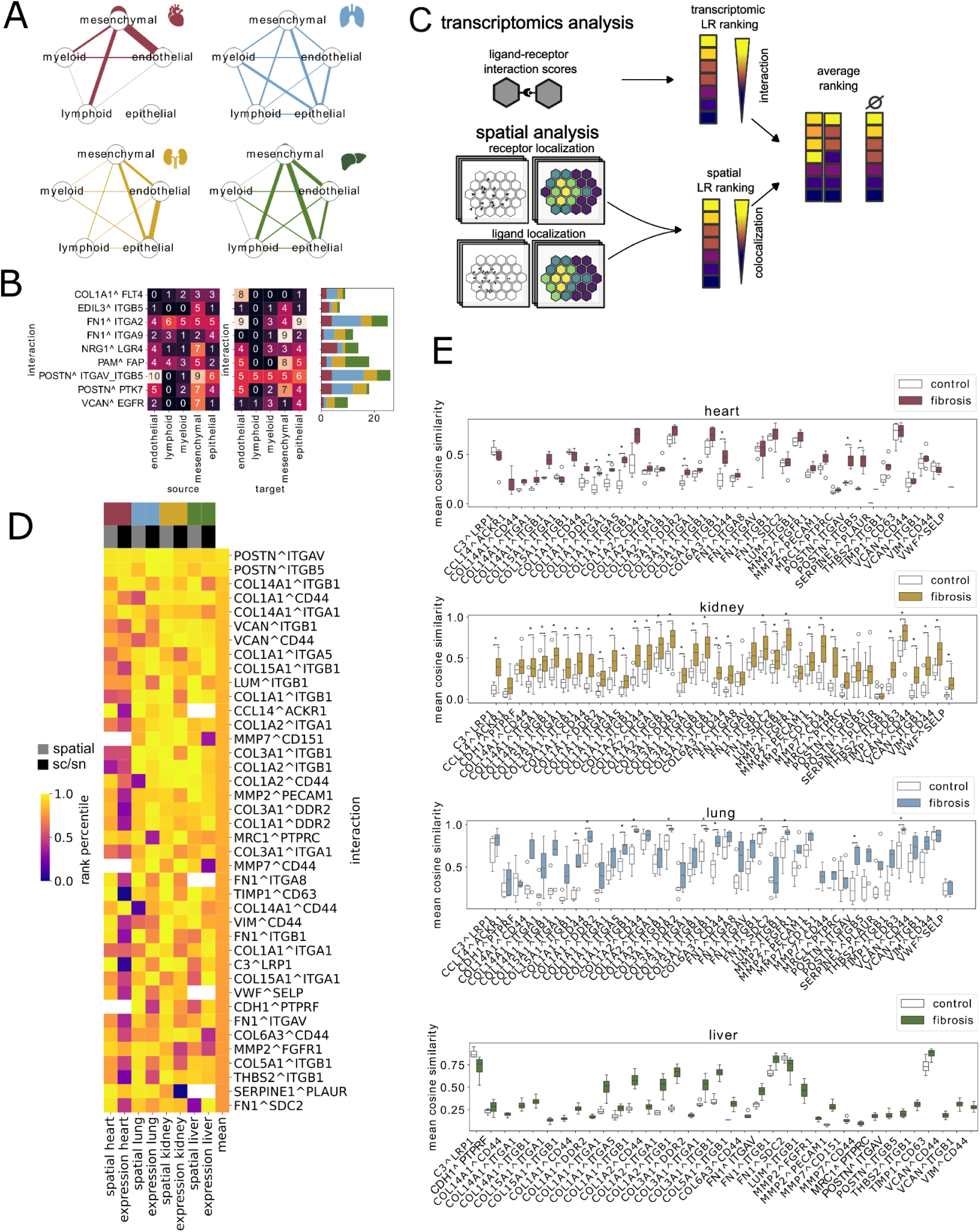
Cell-cell communication in different organs. (A) Upregulated communication (interaction score > 0.5) events in fibrosis between cell types per organ. The thickness of lines reflects the percentage of significant interactions found between the involved cell types. (B) Upregulated communication events (interaction score > 0.5) that are found at least once in all tissues, separated by source (left) and target cell type (right). Bar plot on the right depicts how many times the specific ligand-receptor pair was found in each tissue, regardless of the cell type. (C) Analysis scheme of ligand-receptor analysis. Ligands and receptors with high interaction scores in the transcriptomics analysis and high spatial scar-localization were identified per organ by averaging their ranks of both analyses. (D) Percentile of ligand-receptor pair per organ based on their spatial co-expression rank (grey) or their interaction score rank from communication analysis in the single-cell meta-analysis (black). (E) Mean spatial co-expression, as measured by cosine similarity, of ligand-receptor pairs identified in (D) of fibrosis vs. control patients. Asterisks indicate an adjusted p value < 0.05 based on a two-sided Mann–Whitney U test.

To refine the identification of ligand-receptor interactions relevant to fibrotic pathogenesis, we investigated pairs that were not only transcriptionally upregulated in fibrotic conditions but also exhibited spatial co-localization of expression. Additional spatial transcriptomics datasets were curated to enable this analysis (Table 2). We identified four fibrosis Visium datasets, each representing a distinct organ. In the heart, lung, and liver, the fibrosis and reference groups were each represented by 4-6 samples. In the kidney, more samples were available (13 reference, 40 fibrosis), even though the majority of those consisted of needle biopsies, which leads to a much smaller sample area compared to other datasets (for an example, see Figure S8C).

**Table 2.** Overview of curated spatial transcriptomics datasets.

| Spatial transcriptomics datasets |  |  |  |  |  |
| --- | --- | --- | --- | --- | --- |
| Dataset name | Organ | Samples | Groups | Data type | Study |
| Kuppe_2022 | Heart | 4, 6 | Ref, MI | Visium | Kuppe<sup>25</sup> |
| Mayr_2024 | Lung | 6, 5 | Ref, IPF | Visium | Mayr<sup>84</sup> |
| kpmp_v2 | Kidney | 13, 7, 33 | Ref, H-CKD, DKD | Visium | Lake<sup>85</sup> |
| Andrews_2024 | Liver | 4, 4 | Ref, PSC | Visium | Andrews<sup>86</sup> |

*Ref = reference, MI = Myocardial Infarction (subset to fibrotic-zone samples), IPF = idiopathic pulmonary fibrosis, DKD = diabetic kidney disease, H-CKD = hypertensive chronic kidney disease, PSC = primary sclerosing cholangitis.*

We ranked ligand-receptor pairs per organ by their spatial coexpression in fibrotic samples and by their interaction strength in the transcriptomic data, respectively, regardless of the cell type. We then looked at the ligand-receptor pairs with the highest mean percentile rank across modality and organs (Figure 5C). Similar to the prior analysis, we found many established ligand-receptor pairs, such as periostin (*POSTN*) and fibrillar collagens (*COL1A1, COL3A1, COL5A1*) binding to integrins (*ITGAV*, *ITGA1*, *ITGB5*, *ITGB1*) or discoidin domain receptor (DDR2) (Figure 5D). Furthermore, proteoglycans such as versican (*VCAN*) and lumican (*LUM*) were upregulated in several tissues and were predicted to interact with their receptors based on spatial proximity and gene expression. Some of the top pairs also included ligands or receptors that are not as well-known for their pro-fibrotic roles yet: for instance macrophage mannose receptor 1 (*MRC1*) and receptor-type tyrosine-protein phosphatase C (*PTPRC*, also known as *CD45*). Even though we did not explicitly filter for a spatial colocalization difference between reference and fibrotic samples, but only looked at the spatial colocalization scores of fibrotic samples, we found a general trend of higher colocalization scores of the identified ligand-receptor pairs within fibrotic samples compared to control samples in each organ (Figure 5E). Taken together, we identified the core ligand-receptor interactions in fibrosis by combining our transcriptomic data with spatial co-localization, thereby strengthening the confidence in our findings.

### Scar-specific cell-cell communication

The previous analysis highlighted a broad range of intercellular communication events based on established ligand-receptor interactions. Extracellular matrix components largely dominated these signals, but information regarding their spatial localization, including proximity to fibrotic scar regions, was unavailable. To refine these insights, we next focused on identifying ligands and receptors that are specifically expressed in scarring regions, excluding structural matrix molecules such as collagens and proteoglycans. As an unbiased approach to identify those regions, we calculated the spatial colocalization of all possible ligands and receptors with *COL1A1* in space (see methods), as *COL1A1* was one of the commonly highly expressed genes in fibrosis across all four organs (Figure 4D). Then, we compared the scar-localization of each gene per organ to the organ-consensus scores of disease fibroblasts (Figure 6A). We overall observed a low correlation between scar localization of ligands and receptors and organ effect size in each organ (R in heart = 0.34, liver = 0.38, lung = 0.22, kidney = 0.24), suggesting several cell types and states to be involved in scar-tissue gene expression or a fibrotic gene expression change that goes beyond the scar area (Figure 6B). When comparing the overlap between top ranked genes per organ (upper 20th percentile in gene regulation and colocalization), we observed 7 genes that were identified in 3 out of 4 organs (*VIM, TIMP1, FSTL1, CCN2, ANXA2, FBN1, FN1, SPON1, THBS2*), and 2 genes (*TIMP2, MRC2*) that were identified in all four organs (Figure 6C). Finally, we sorted all investigated genes by their average rank across all eight categories (1 spatial coexpression and 1 organ-consensus score per organ) (Figure 6D). Our top hit was Metalloproteinase inhibitor 1 (*TIMP1*), whose spatial co-expression with *COL1A1* was detected in fibrotic samples from all four organs (Figure S8A-D) and whose organ-effect size was consistently positive across organs in disease fibroblasts (Figure S8E). *TIMP1* is a well-known profibrotic gene that has been shown to have an effect in several organs^87–89^. Further top hits on our list included other well-known and –explored pro-fibrotic genes such as *THBS2*^90–93^, *POSTN*^76–79^, and *TNC*^76,94–96^. Recent studies identified inhibition of TNC as an effective therapeutic strategy in a murine model of systemic sclerosis^97^. Our findings further suggest that targeting TNC may hold broader therapeutic potential across a spectrum of fibrotic diseases.

**Figure 6.**
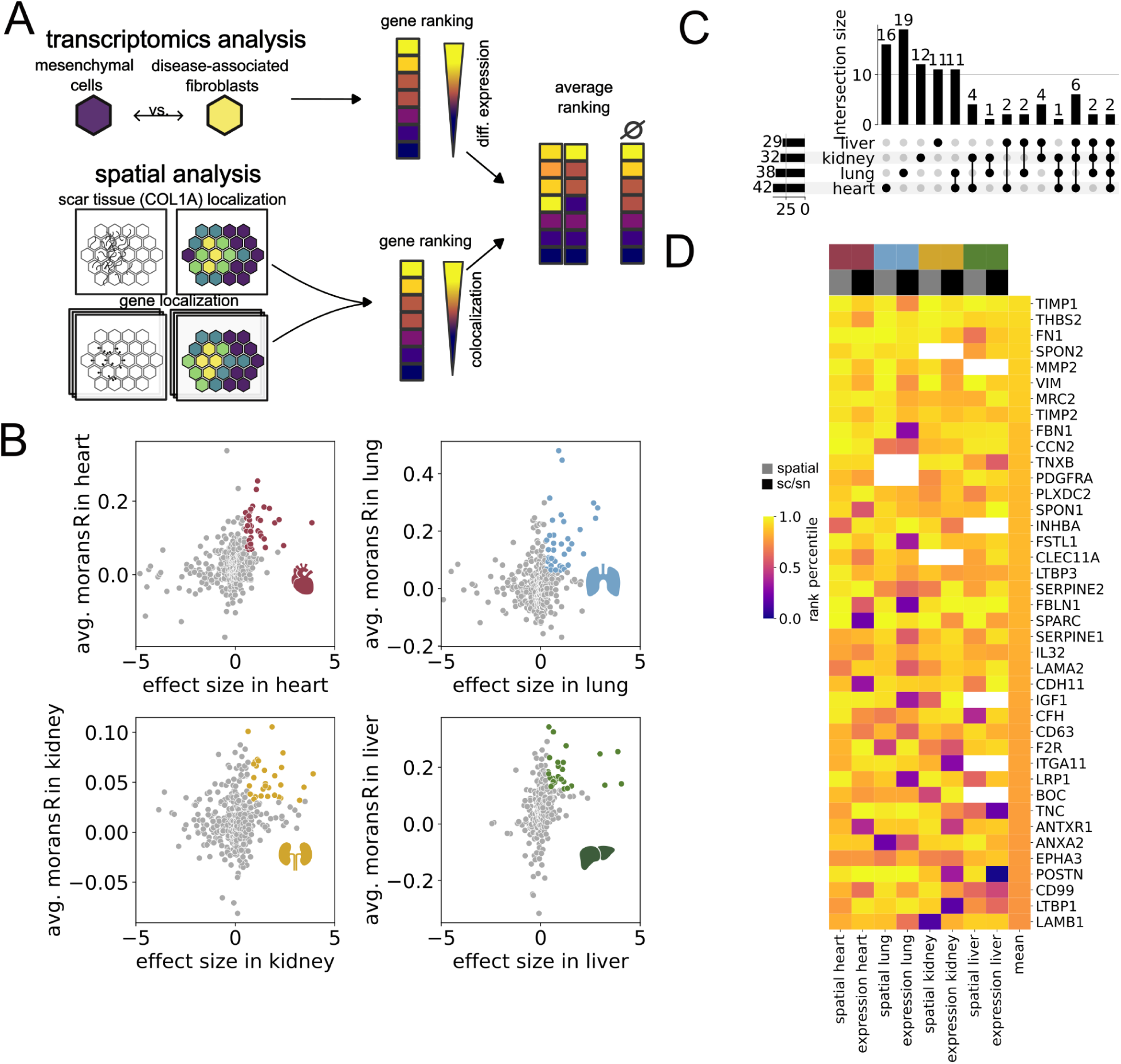
Identification of fibrotic targets. (A) Analysis scheme of target identification. Ligands and receptors with high differential gene expression in disease-associated fibroblasts and high spatial scar localization were identified per organ by averaging their ranks from both analyses. (B) Avg. Moran’s R and organ effect size of disease-associated fibroblasts for all tested genes. Marked genes reflect the genes in the upper 20th percentile. (C) Overlap of marked genes from (B) across tissues. (D) Ranked heatmap depicting the percentile of average Moran’s R of fibrotic visium slides & organ-effect sizes of disease-associated fibroblasts per organ and gene.

## Discussion

In this study, we curated a large collection of fibrosis-related datasets from 20 single-cell studies. This comprehensive analysis spanned multiple levels, from global comparisons of cellular composition and predictive disease signatures to detailed insights into individual genes, transcription factors, and intercellular communication events that may be involved in fibrotic remodeling.

While we found strong differences in cell type composition changes between tissues and, to a lesser extent, between studies, we observed a common pattern of fibrosis gene regulation across most tissues. Liver tissue represented an exception, displaying the greatest study-to-study variability. We attribute this, in part, to technical heterogeneity across liver datasets. The weaker agreement across liver studies may also reflect greater biological variability, which could potentially be caused by the pronounced capability for regeneration by the liver. This regenerative potential likely results in a broader range of fibrotic phenotypes within patients diagnosed with the same condition, spanning minimally fibrotic tissue, highly cirrhotic states, and partially healed scars. At the opposite end of the spectrum, heart and lung datasets primarily included end-stage disease samples, which may represent a more homogeneous disease population and thus exhibit stronger cross-study concordance.

Cross-tissue comparisons revealed strong overlap in fibrotic gene regulation between heart and lung tissue in mesenchymal and endothelial cells. Interestingly, even though myeloid cells showed high transcriptomic similarities within lung and heart tissues, they showed low overlap in gene regulation between the two tissues. To identify which genes show the highest agreement in fibrotic regulation, we focused on genes that are most consistently up– or downregulated across datasets. As expected, we recovered many known pro-fibrotic genes in our analysis. However, we also identified numerous genes that have either not been previously implicated in fibrotic disease or have been studied only within a single tissue context. For instance, we identified fibroblast growth factor 14 (*FGF14*) and DBH-like monooxygenase protein 1 (*MOXD1*) as commonly upregulated in mesenchymal cells across organs. *FGF14* was reported to be a marker of a pro-fibrotic fibroblast subset in the kidney^29^ and lung^98^, but has not been functionally described or reported in liver or heart fibrosis. On the other hand, *MOXD1* has been previously shown to be upregulated in all organs^48,49,99,100^. However, its experimental downregulation was tested only in the liver^48^ and kidney^49^, but not in the other organs to our knowledge. Our results suggest that inhibition of *MOXD1* could offer broader therapeutic benefits across diverse fibrotic conditions. We also explored cell-state-specific gene expression and the spatial localization of ligands and receptors. Here, we identified Tenascin C as specifically upregulated in disease-associated fibroblasts in scar regions. While Tenascin C inhibition has been shown to be an effective therapeutic strategy in a murine model of systemic sclerosis^97^, our findings further suggest that targeting it may hold broader therapeutic potential across a spectrum of fibrotic diseases. Thus, by integrating multiple studies per organ and extending the comparison across organs, our meta-analysis enables the (re-)discovery of genes that have been investigated only in a specific biological setting or not at all.

Our study has several limitations. First, substantial technical differences across datasets, including variation in sequencing modality (e.g., single-cell RNA sequencing vs. single-nucleus RNA sequencing), as well as differences in sampling and sequencing methods, likely reduce overlap between studies and organs and may obscure genes that play important roles in fibrotic processes. Second, the limited availability of patient metadata leaves many aspects unresolved, including the exact diagnosis, disease severity, tissue sampling location, and the extent of fibrosis. If these aspects were better documented, they could be accounted for in the analysis and could allow a clearer distinction of physiological from pathophysiological fibrotic processes. Third, we treated all disease etiologies collectively under the term “fibrosis”. However, the degree of fibrotic remodeling likely varies between conditions, and the dataset remains imbalanced in terms of sample representation across organs. Lastly, with exception to the mesenchymal cell population, our analysis primarily focused on broad cell type categories, even though some studies have highlighted the disease-relevance of specific cell states. Data generated using standardized protocols across different fibrotic disease etiologies would facilitate more detailed characterization in this regard. Nevertheless, the recurrence of shared gene regulation patterns across studies and tissues using different analysis methods strongly supports the existence of common pro-fibrotic and anti-fibrotic molecular programs.

Beyond the highlighted examples of shared molecular features across tissues, we identified many additional genes and patterns of interest within each cell type. To facilitate further exploration, all results from this analysis are publicly available as an interactive resource at https://organfibrosis.saezlab.org/ (Figure S9). Users can query analysis results from our differential gene expression analyses for each major cell type and disease associated fibroblasts. In addition, ligand-receptor interaction scores, their spatial colocalization, and scar colocalization scores are available. This platform is intended to support hypothesis generation and to enable researchers to examine expression trends or potential regulatory mechanisms relevant to their genes of interest.

Taken together, our study provides a large, integrative resource that captures both organ-specific and shared gene programs of fibrosis, offering a framework that can be continuously expanded as new single-cell and spatial datasets emerge.

## Methods

### Processing of single-cell and single-nucleus data

Datasets were downloaded from their publicly available repositories and, if not already the case, transformed into anndata objects. Available metadata nomenclature was harmonized. For each dataset, cells with fewer than 200 genes or 500 counts, or more than 40000 counts, were removed. Genes present in fewer than three cells were filtered. Furthermore, cells with more than 25% mitochondrial gene content were removed, except for the datasets Abedini_2024_sc and Lake_2023_sc, where the mitochondrial gene cutoff was increased to 35% due to exceptionally high mitochondrial gene counts in kidney scRNA data.

### Cell type harmonization

Cell type labels from the authors were utilized whenever they fit into our defined categories of endothelial, lymphoid, myeloid, epithelial, or mesenchymal cells. A list of cell types and how they were grouped can be found in Suppl. Table 1. For two studies, namely Muto_2022^29^ and Wilson_2022^31^, not all cells could be clearly allocated to those cell type categories. Therefore, the cells were reannotated using scVI and scANVI from scvi-tools^101,102^ (v1.1.6.post2) to learn batch-corrected latent representations and transfer cell type annotations from Lake_2023_sn^26^ to the respective dataset. To build a reference model, the Lake_2023_sn dataset was normalized to 10000 counts per cell, log-transformed, and restricted to highly variable genes selected in a batch-aware manner. A scVI model was trained on raw counts to learn a low-dimensional latent space. The model was subsequently transformed into a scANVI model. For label transfer, the trained scANVI model was used as a reference, and each study was treated as a query dataset after harmonizing gene features and masking existing annotations. Cell type predictions were then generated for the query data using the fine-tuned model.

Cell type marker genes were identified by contrasting pseudobulks of one cell type against the remaining cell types. Pseudobulks were calculated for each sample by summing the counts of all cells in a given cell group. Their marker gene expression was assessed using edgeR (v3.40.0)^103–106^ with default gene filtering settings. Marker gene overlap between studies was evaluated by calculating the Jaccard Index for all marker genes across studies and cell types, with an FDR < 0.01 and logFC > 2.

### Quantification of transcriptional changes

To assess disease-associated transcriptional shifts within each cell type, we applied *scDist* (v1.1.2)^107^ to estimate multivariate transcriptional distances between fibrotic and control samples. Single-cell count matrices were filtered to retain annotated cell types present in both conditions. Raw counts were normalized using Seurat (v5.1.0)^108^ SCTransform function. The *scDist* model was then fitted using disease condition as a fixed effect and sample as a random effect, stratified by cell type. The method estimates a global transcriptional distance per cell type and identifies the genes that contribute most strongly to this disease-associated shift. Studies with an estimated effect size of 0 in more than 2 cell types were excluded from further analysis.

### Compositional data analysis

Cell type proportion changes between fibrosis and control samples were analyzed separately for each organ. Proportions were first transformed using the centered log-ratio (CLR) transformation. For each cell type within each study, Welch’s *t*-tests were performed to compare CLR-transformed proportions between conditions, followed by Benjamini-Hochberg correction for multiple testing. In addition, linear mixed-effects models (*value* ∼ *condition* + (1|study)) were fitted per cell type to estimate condition effects while accounting for study-specific variability. Effect estimates, *t*-statistics, and adjusted *p*-values were visualized as barplots and heatmaps to summarize organ-specific changes in cell type abundance.

### Multicellular factor analysis

Molecular variability across cell types and patient samples was summarized as multicellular programs, latent factors capturing shared gene expression patterns across cell types across all studies from one organ. Each factor can be associated with a molecular phenotype that might be related to clinical or biological annotations.

We estimated 15 factors per organ using Multicellular Factor Analysis using *MOFA2* (v1.9.2)^36,109^. Pseudobulk expression profiles were included only for cell types represented by at least 25 cells and present in ≥40% of patient samples with more than 50 detected genes. Genes with fewer than 100 counts in any sample or detected in <40% of samples were excluded. Expression values were normalized with trimmed mean of M-values (TMM) normalization (*edgeR* v3.40.0) with a scale factor of 1,000,000 and subsequently log-transformed. Marker genes of other cell types were removed to minimize contamination, and samples with <97% gene coverage were excluded to avoid coverage-related artifacts. Each study was treated as a separate group in the MOFA model.

The association between factor scores and disease status (fibrosis vs. reference) was assessed using ANOVA with Benjamini-Hochberg correction (FDR ≤ 0.05).

### Differential gene expression (meta-)analysis

Differential gene expression analysis was performed between fibrosis and reference samples independently for each cell type and study using PyDESeq2 (v0.5.0)^110^. Pseudobulk expression matrices were filtered using expression thresholds (≥ 10 counts per gene in 70% of samples and ≥15 counts per gene across all samples), and analyses were restricted to cell types with at least three samples per condition. Normalization, dispersion estimation, and log₂ fold-change calculations were performed using the DESeq2 framework, with LFC shrinkage applied to stabilize effect estimates.

For the meta-anlaysis across datasets, resulting gene-wise statistics were aggregated and subjected to a random-effects meta-analysis for each cell type using the statsmodels (v0.14.4) combine_effects function with the “iterated” method to iteratively estimate the between-study variance^111,112^. As model inputs, the log₂ fold change and its squared standard error were used for each gene. Gene-wise meta-analyses were first performed within each organ to derive organ-specific summary effect sizes, followed by a second meta-analysis across organs to identify consistent transcriptional signatures shared between tissues. For both models, genes with differential gene expression results (for the organ-specific model) or with effect sizes (for the cross-organ model) in fewer than two studies and organs, respectively, were skipped. Genes with an organ effect size > 0.5 and a lower confidence interval > 0 were classified as upregulated organ-consensus genes; genes with a combined effect size < –0.5 and an upper confidence interval < 0 were classified as downregulated organ-consensus genes.

### Functional interpretation of common gene programs

To functionally interpret the meta-analytic differential expression results, we inferred pathways and transcription factor activities using the *decoupler*^113^ framework (v1.9.0). Organ-specific meta-analysis effect sizes normalized by their standard deviation were used as input. Gene set resources were compiled from MSigDB^114,115^ (Gene Ontology biological process collections) and CollecTRI^50^ transcription factor regulons. For each gene set collection, univariate linear model (ULM) enrichment analyses were performed to estimate activity scores and corresponding *p*-values per organ and gene set. To identify commonly regulated gene programs across organs, gene sets were first ranked by the mean ULM enrichment statistic across organs. To ensure biological consistency, only gene sets whose enrichment scores were strictly positive or strictly negative in all organs were retained and visualized.

### Patient classification models

To assess the concordance of transcriptional regulation across studies, we conducted a cross-study disease-prediction analysis. For each cell type, we used lists of differentially expressed genes (DEGs) identified in one study (“training study”) to predict the disease status of all samples in another study (“test study”) using their pseudobulk expression profiles. The top 500 DEGs were selected according to absolute t-statistics. Pseudobulk expression data were normalized, log-transformed, and scaled prior to prediction. For each sample, the dot product between DEG t-values from the training study and normalized gene expression values from the test study was computed as a disease activation score. Predictive performance was quantified using the area under the receiver operating characteristic curve (AUROC) comparing fibrosis versus control samples. Cross-study and cross-organ AUROC values were later aggregated by computing organ-level median AUROC values to quantify the similarity of regulated gene programs across tissues.

### Integration of mesenchymal cells and disease-associated fibroblast definition

To identify disease-associated mesenchymal cells, mesenchymal cells were subsetted from the datasets based on cell type annotations. To identify highly variable genes (HVGs) across organs, they were first selected per study and then combined with the results of other studies from the same organ. First, samples were filtered to those containing at least ten mesenchymal cells. Cells expressing fewer than 200 genes and genes detected in fewer than ten cells were removed. Expression values were normalized to 10,000 counts per cell and log-transformed. HVGs were then identified per sample using scanpy’s (v1.10.4)^116,117^ highly variable gene function with thresholds of min_mean=0.0125, max_mean=3, and min_disp=0.5, and the proportion of samples in which each gene was selected as HVG was computed. The final HVG set for each organ included genes reported as highly variable in the majority of samples. For each organ, the resulting HVG matrices were used to train scVI models (v1.1.6.post2)^101,118^ to obtain latent representations of mesenchymal transcriptional states. Raw counts were modeled using a negative binomial likelihood, with the study as the batch covariate and modality and sequencing technique as categorical covariates. Continuous covariates included sequencing depth, the number of detected genes, and mitochondrial and ribosomal gene proportions. Each model was trained for 50 epochs with 30 latent dimensions, using deterministic random seeds to ensure reproducibility. The learned scVI embeddings were subsequently clustered with the Leiden algorithm at several resolutions. Resolutions resulting in 5-7 clusters were manually selected. For each organ, compositional data analysis was performed on the resulting clusters, as described in the compositional data analysis section, using clusters instead of cell types. Considering the compositional changes of clusters across studies, together with myofibroblast marker gene expression, and author annotations of individual studies (not available for all studies), led to the annotation of respective clusters as “disease-associated fibroblast” clusters. The contrastive analysis described above was repeated, comparing disease-associated fibroblasts with non-disease-associated mesenchymal cells rather than fibrosis with non-fibrosis samples. Differential gene expression results were again summarized into organ and cross-organ effect sizes.

### Cell-cell communication analyses

To infer organ-specific cell-cell communication patterns from gene expression meta-analysis results, we integrated ligand-receptor (LR) resources with model-derived effect sizes. For each organ and cell type, ligand and receptor effect sizes and standard deviations were mapped to consensus LR pairs from the LIANA^119^ consensus resource (v1.5.0). All pairwise source-target cell type combinations were evaluated to assemble organ-level communication networks. For each LR pair, ligand and receptor organ effect sizes were averaged to compute interaction scores, and complex subunits were collapsed using a minimum-expression policy to handle multi-subunit interactions.

### Processing of spatial transcriptomics data

For each organ, one spatial transcriptomics dataset was curated. Spots with ≤ 400 gene counts or ≤ 5 cells were filtered. When the data was not already normalized, each spot was normalized by total counts over all genes with a target sum of 10,000 and subsequently log-transformed. No cell type annotation was used in the subsequent data analysis.

### Spatial cell-cell communication analysis

To combine information from spatial transcriptomic datasets and the single-cell meta-analysis, ligand-receptor pairs were ranked based on their interaction score from the transcriptomic cell-cell communication analysis within each organ. To estimate their spatial colocalization in each organ, we calculated the bivariate Moran’s R metric^120,121^ implemented in LIANA+^119^ v1.5.0 per Visium slide, and the average Morans’s R metric for fibrotic samples was ranked for each organ and ligand-receptor pair. Finally, all ligand-receptor pairs were sorted by their average rank across both modalities and tissues.

### Target identification analysis

To identify potential targets to explore for therapeutic exploration, we combined information from the single-cell meta-analysis with spatial datasets. We defined targets of interest as ligands and receptors present in the LIANA+^119^ consensus resource, but excluded Collagens and Proteoglycans as defined in the NABA_COLLAGENS and NABA_PROTEOGLYCANS^122^ gene sets. Then, the coexpression of each ligand and receptor with *COL1A1* was estimated with the bivariate Moran’s R metric implemented in LIANA+^119^ v1.5.0 per Visium slide. The mean Moran’s R metric^120,121^ for fibrotic samples per tissue was ranked for each organ and gene. Similarly, the same gene list was ranked by organ effect size. Finally, all genes were sorted by their average rank across both modalities and tissues.

## Code & data availability

The code used for data analysis was formatted into a snakemake^123^ pipeline and is publicly accessible at https://github.com/saezlab/organfibrosis. All datasets used were publicly available and can be found in the respective publications listed in Tables 1 & 2. To facilitate further exploration of the data, reprocessed data is available for download on zenodo (DOI: 10.5281/zenodo.18432221) and can be visualized and downloaded at https://organfibrosis.saezlab.org/.

## Acknowledgements

This work was supported by the German Federal Ministry of Research, Technology, and Space (BMBFTR) with the project 01EJ2201B “Curefib”. Furthermore, the authors acknowledge support by the state of Baden-Württemberg through bwHPC and the German Research Foundation (DFG) through grant INST 35/1597-1 FUGG. The results presented here are in part based upon data generated by the Kidney Precision Medicine Project (KPMP). The KPMP is supported by the National Institute of Diabetes and Digestive and Kidney Diseases (NIDDK) through the following grants: U01DK133081, U01DK133091, U01DK133092, U01DK133093, U01DK133095, U01DK133097, U01DK114866, U01DK114908, U01DK133090, U01DK133113, U01DK133766, U01DK133768, U01DK114907, U01DK114920, U01DK114923, U01DK114933, U24DK114886, UH3DK114926, UH3DK114861, UH3DK114915, and UH3DK114937. We gratefully acknowledge the essential contributions of our patient participants and the support of the American public through their tax dollars. The authors also acknowledge the University of Michigan Medical School Central Biorepository (RRID:SCR_026845) for providing biospecimen storage, management, and distribution services in support of the research reported in this publication/grant application/presentation. The content is solely the responsibility of the authors and does not necessarily represent the official views of the National Institutes of Health. AI-assisted tools were used to improve the clarity and style of the manuscript; all scientific content and interpretations were authored by the authors. We thank Philipp Sven Lars Schäfer, Chiara Schiller, Charlotte Boys, Robin Falleger, Nils Englert, and Christoph Mahler for their discussions and input on this project.

## Authors contributions

The project was conceived by LK, RORF, and JSR. The analysis was developed and performed by LK with input from JDL and MK, and supervision by RORF and JSR. The website resource was developed by GK. The manuscript was drafted by LK, edited by RORF, and JSR, and jointly revised by all co-authors.

## Supplementary Materials

**Supplementary Figure 1.**
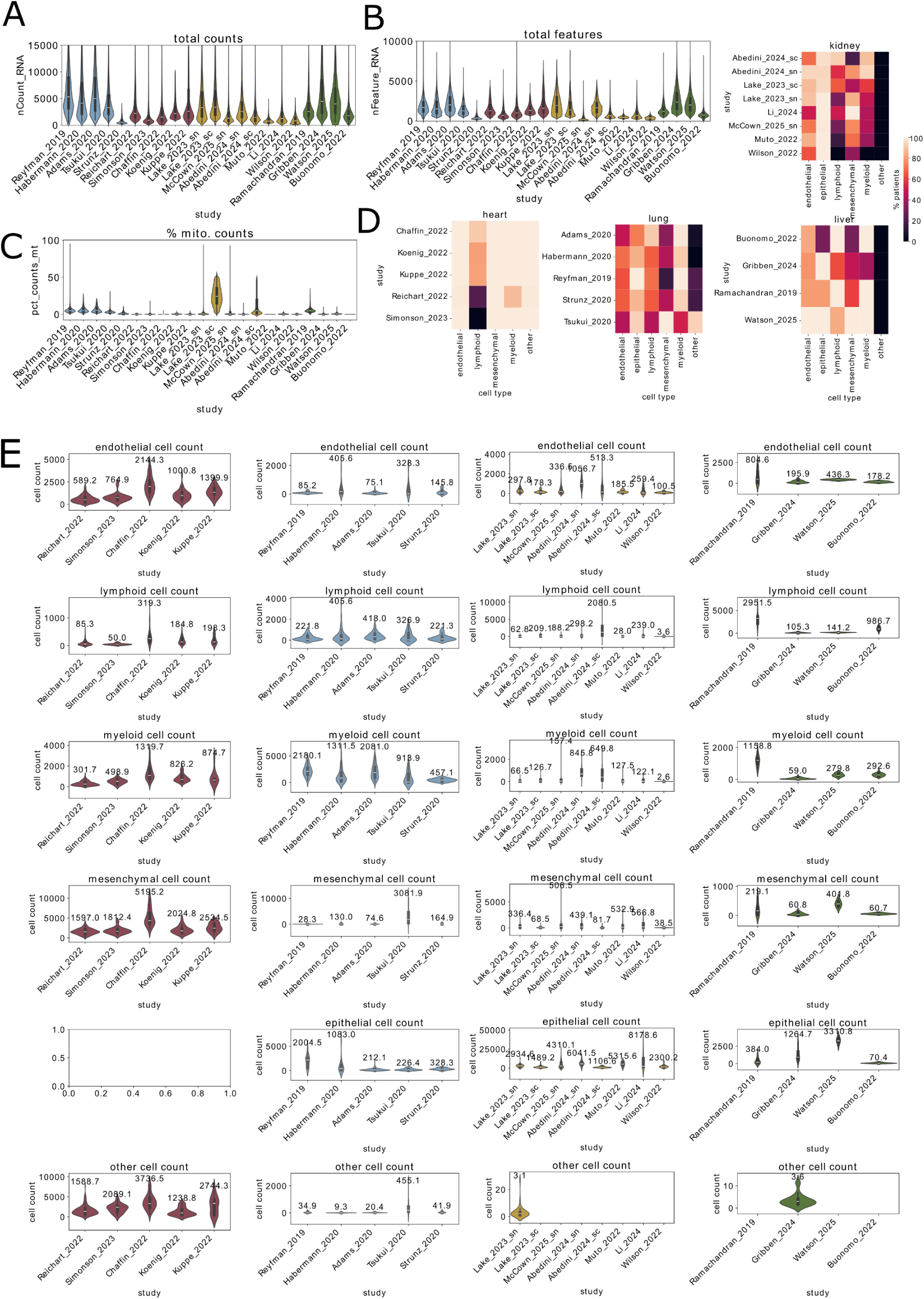
Quality measures. (A) Distribution of total counts per cell, separated by study. Each study is colored by their tissue of origin (blue = lung, red = heart, yellow = kidney, green = liver). (B) Distribution of total features per cell, separated by study. (C) Distribution of percentage of mitochondrial gene counts per cell, separated by study. (D) Percentage of samples with more than 50 cells of a certain cell type, separated by study (from left to right: heart, lung, liver; kidney on top). (D) Cell counts per sample, separated by study & cell type (one cell type per row) and tissue (one tissue per column – from left to right: heart, lung, kidney, liver). The number above each violin denotes the mean per study.

**Supplementary Figure 2.**
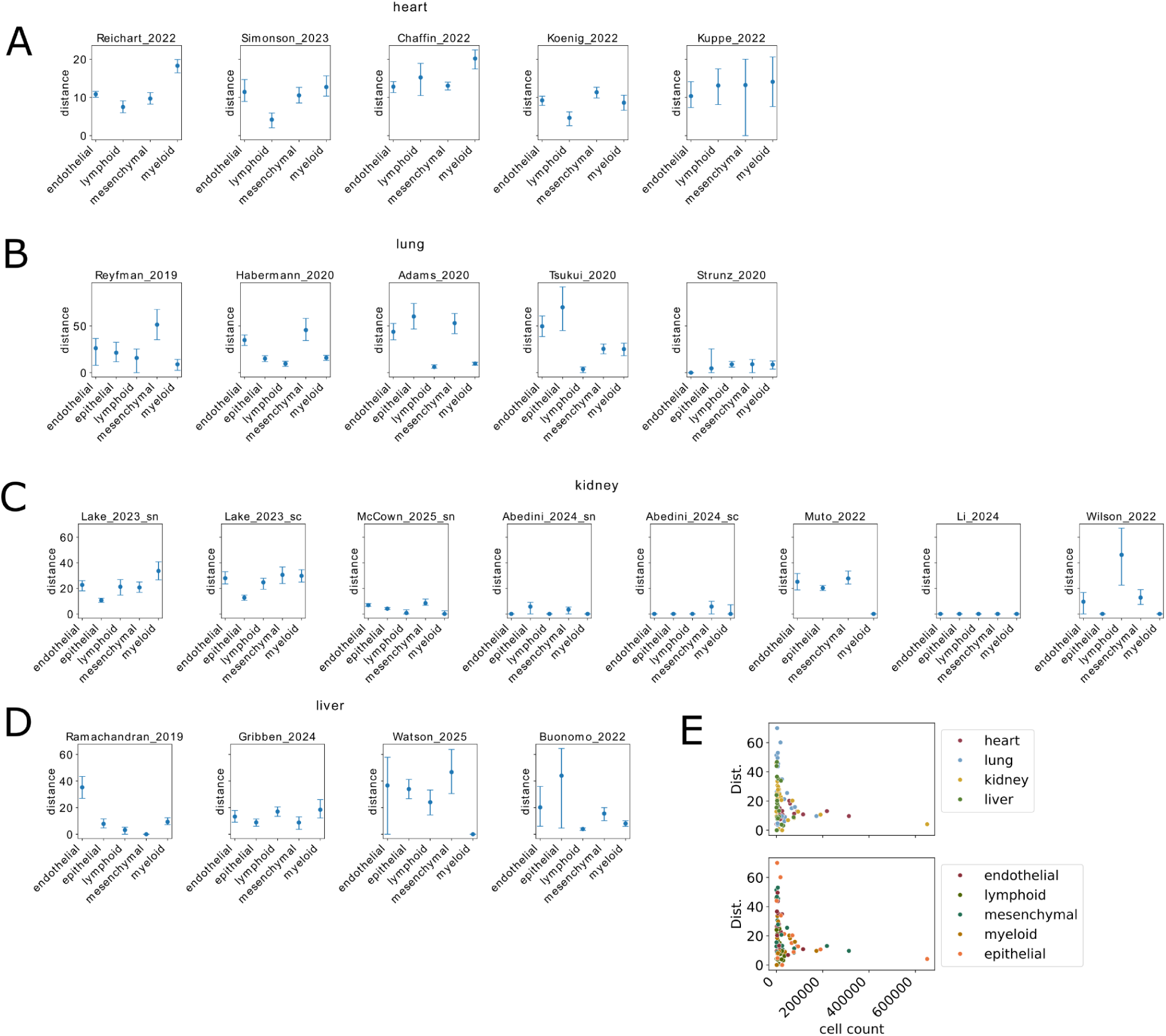
Transcriptomic distance estimations. (A-D) Transcriptomic distances between reference and fibrosis patients estimated by scDist per study and cell type in heart (A), lung (B), kidney (C), or liver (D). (E) Relationship between cell count per study and cell type and estimated transcriptomic distances.

**Supplementary Figure 3.**
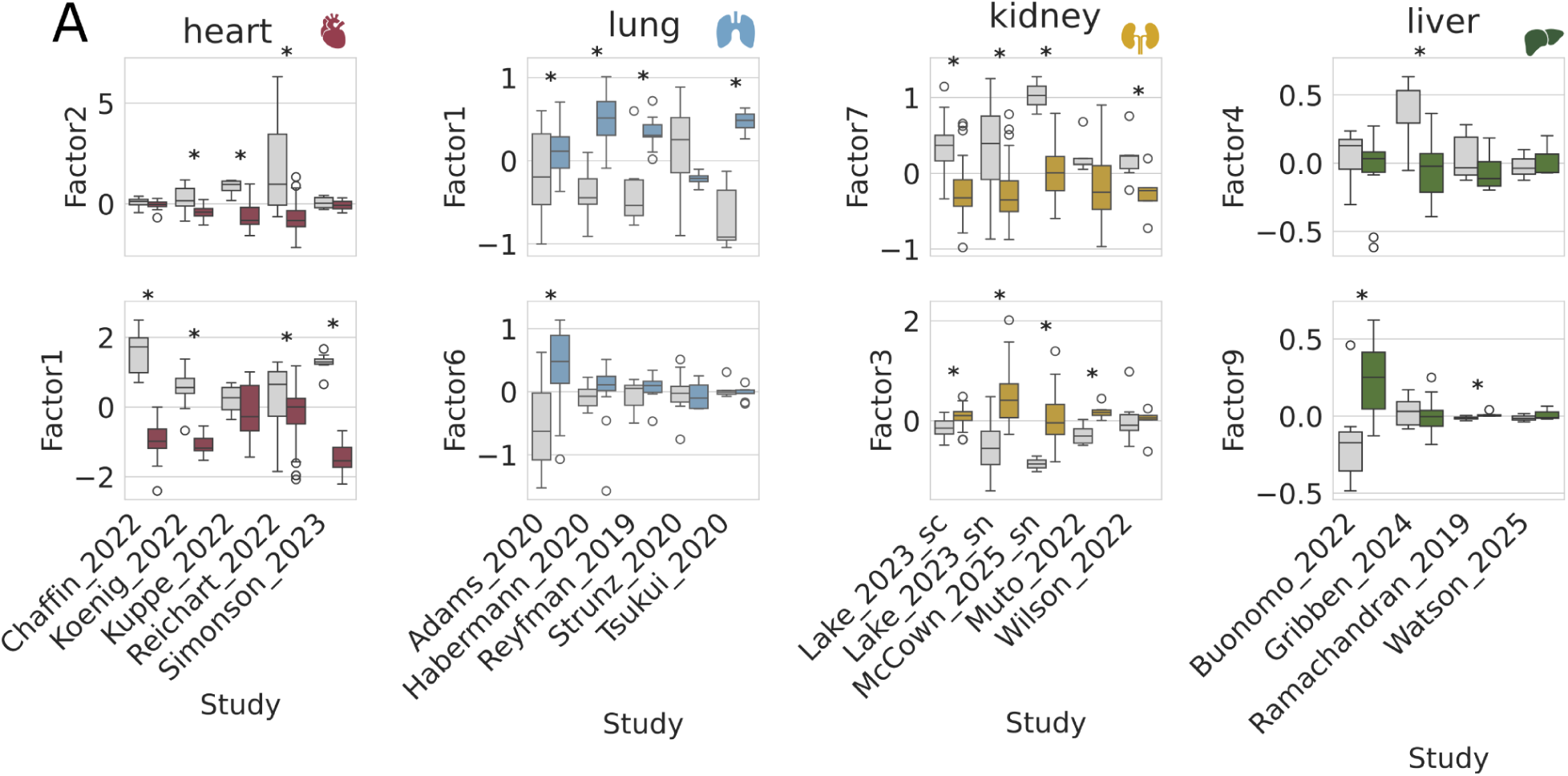
Latent factors associating to fibrosis. (A) Factor loadings per study and patient group of the two factors that separate fibrotic samples best from non-fibrotic samples. The association between factor scores and disease status (fibrosis vs. reference) was assessed using ANOVA with Benjamini-Hochberg correction (* = p.adj. < 0.05). From left to right: heart, lung, kidney, liver.

**Supplementary Figure 4.**
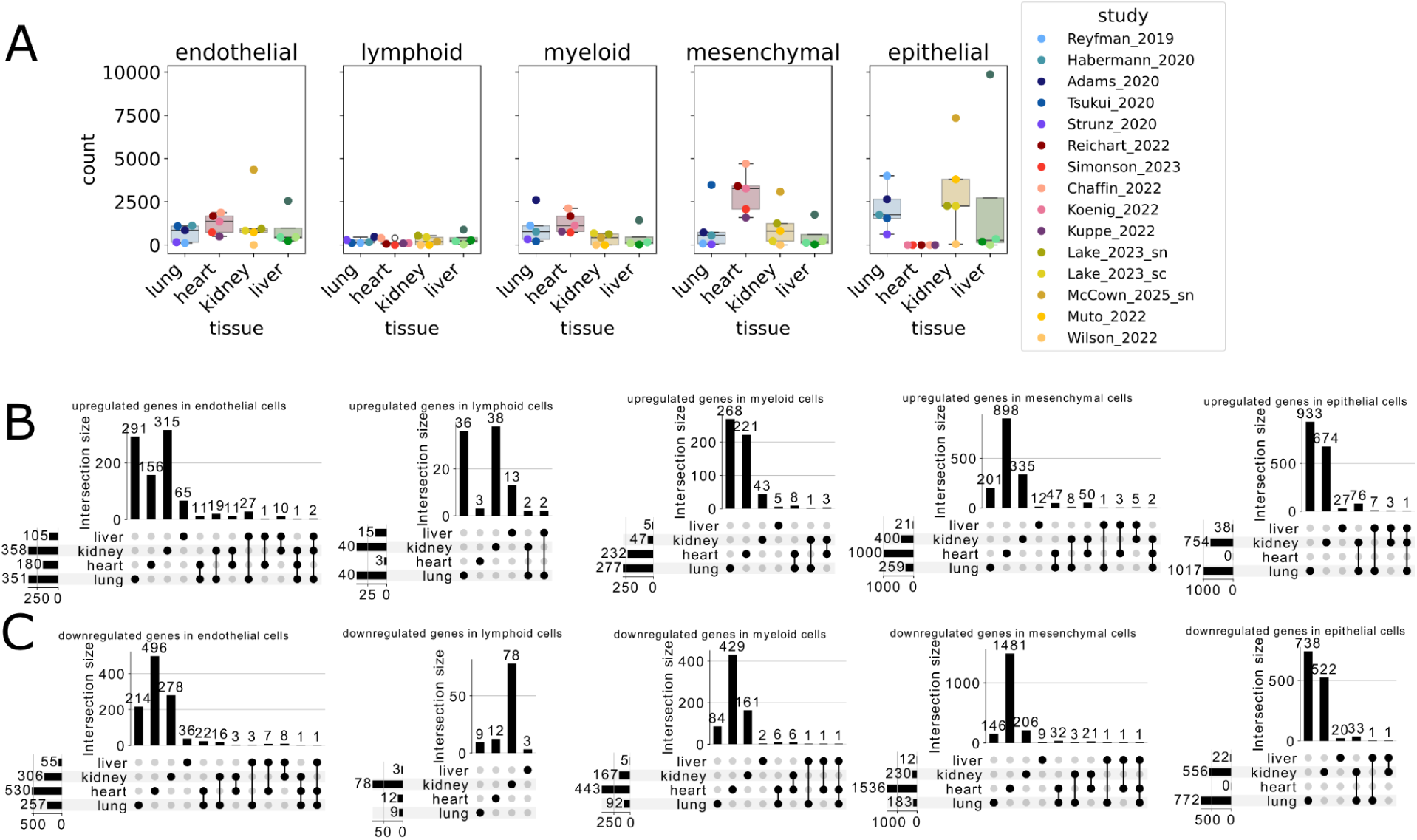
Differential gene expression comparison. (A) Absolute number of differentially expressed genes (log2FC > 0.5 or log2FC < –0.5, p.adj. < 0.05) per study and cell type. The studies are separated by organ they belong to (blue: lung, red: heart, yellow: kidney, green: liver). (B-C) Overlap of upregulated (B) and down-regulated (C) organ consensus genes across tissues. From left to right: Endothelial cells, lymphoid cells, myeloid cells, mesenchymal cells, epithelial cells. A gene was defined as an upergulated organ consensus gene with a combined effect size > 0.5 and lower confidence interval > 0 or as downregulated with a combined effect size < –0.5 and a upper confidence interval of < 0.

**Supplementary Figure 5.**
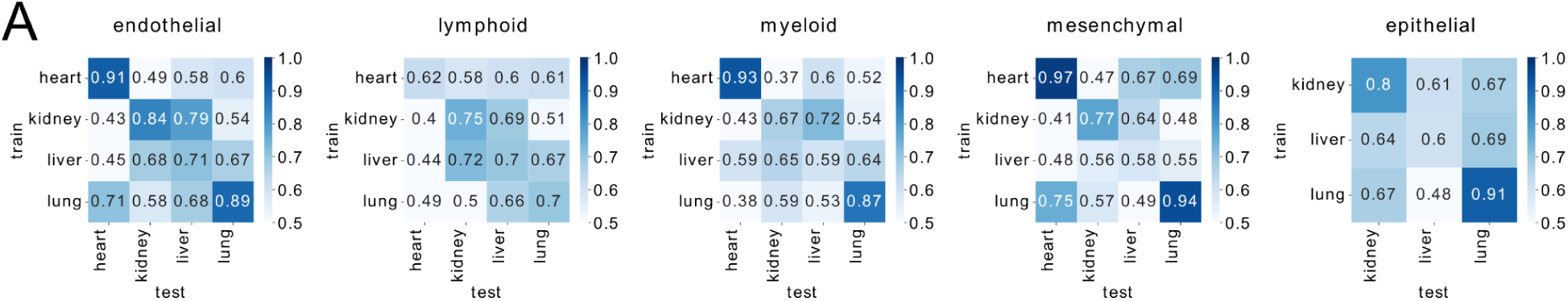
Disease score predictions without organ-consensus genes. Median AUROC values from disease score predictions across tissues per cell type, excluding organ-consensus genes (effect size of > 0.5 & lower confidence interval > 0 / effect size < –0.5 & upper confidence interval of < 0) from both tissues that are compared.

**Supplementary Figure 6.**
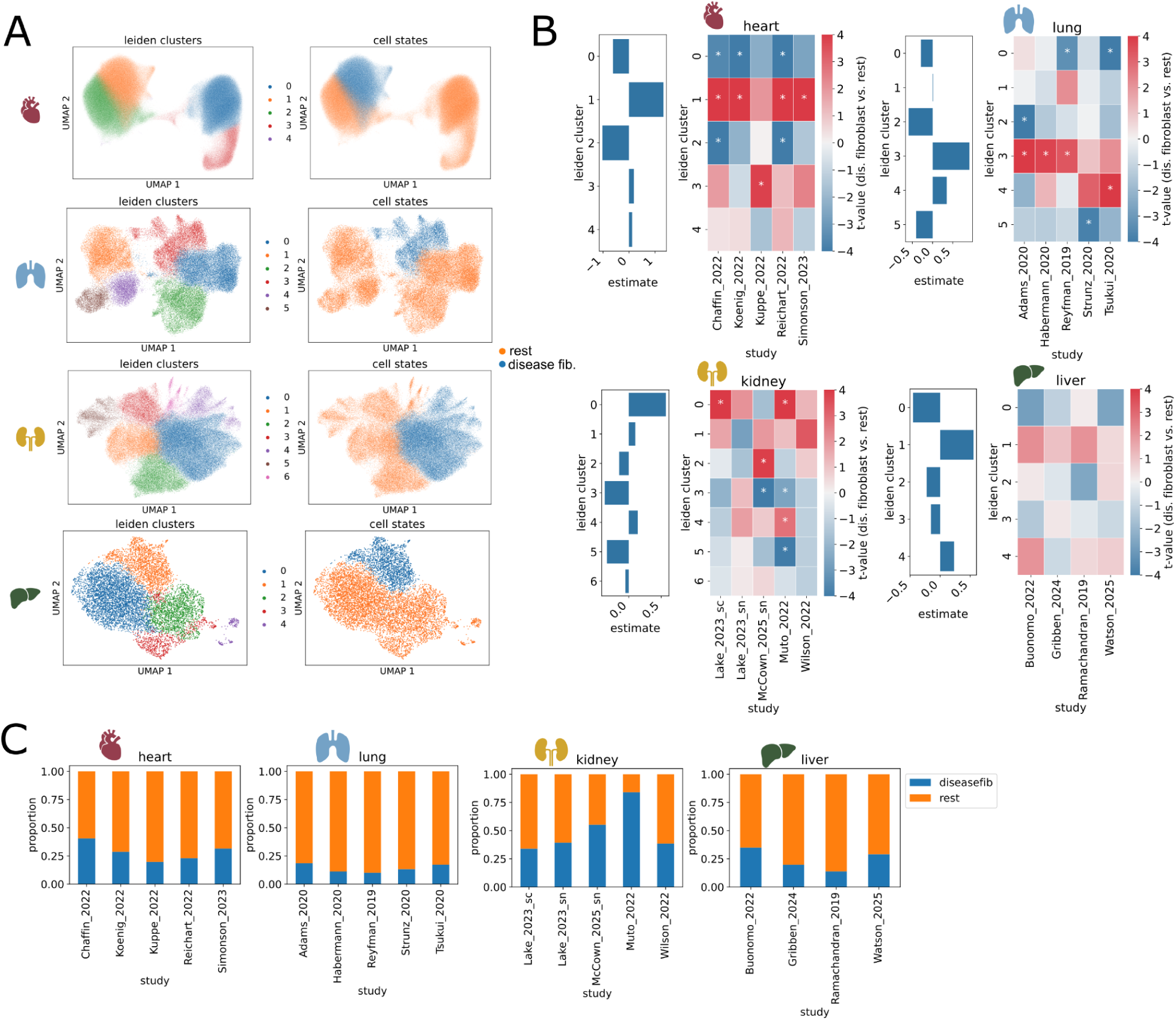
Disease-associated fibroblast cell state identification. (A) UMAP of all mesenchymal cells colored by Leiden clusters (left) and acquired cell state annotation (right) per scvi organ integration (from top to bottom: heart, lung, kidney, liver). (B) Differential compositional analysis per leiden cluster: T-statistics (heatmap) and estimate (bars) of difference between fibrotic and non-fibrotic samples using t-tests and linear mixed-models, respectively. Stars denote an adj. p-value < 0.05. (C) Percentage of mesenchymal cell labels as disease-associated fibroblast (blue) and “rest” per study (from left to right: heart, lung, kidney, liver).

**Supplementary Figure 7.**
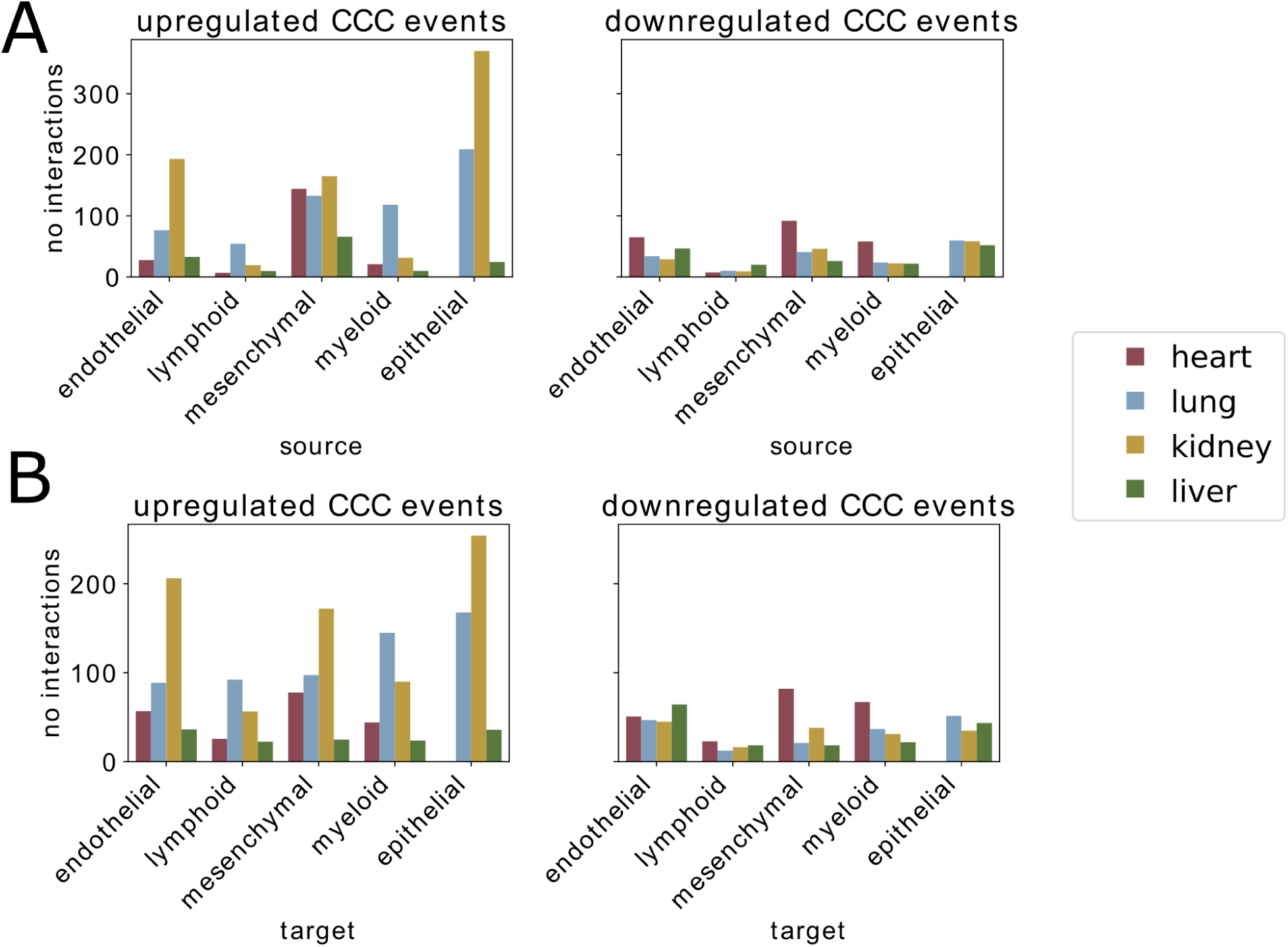
Absolute number of communication events per organ. (A) Absolute numer of upregulated (left) and downregulated (right) communication interactions per organ and cell type, separated by cell type expressing the ligand. (B) Absolute numer of upregulated (left) and downregulated (right) communication interactions per organ and cell type, separated by cell type expressing the target.

**Supplementary Figure 8.**
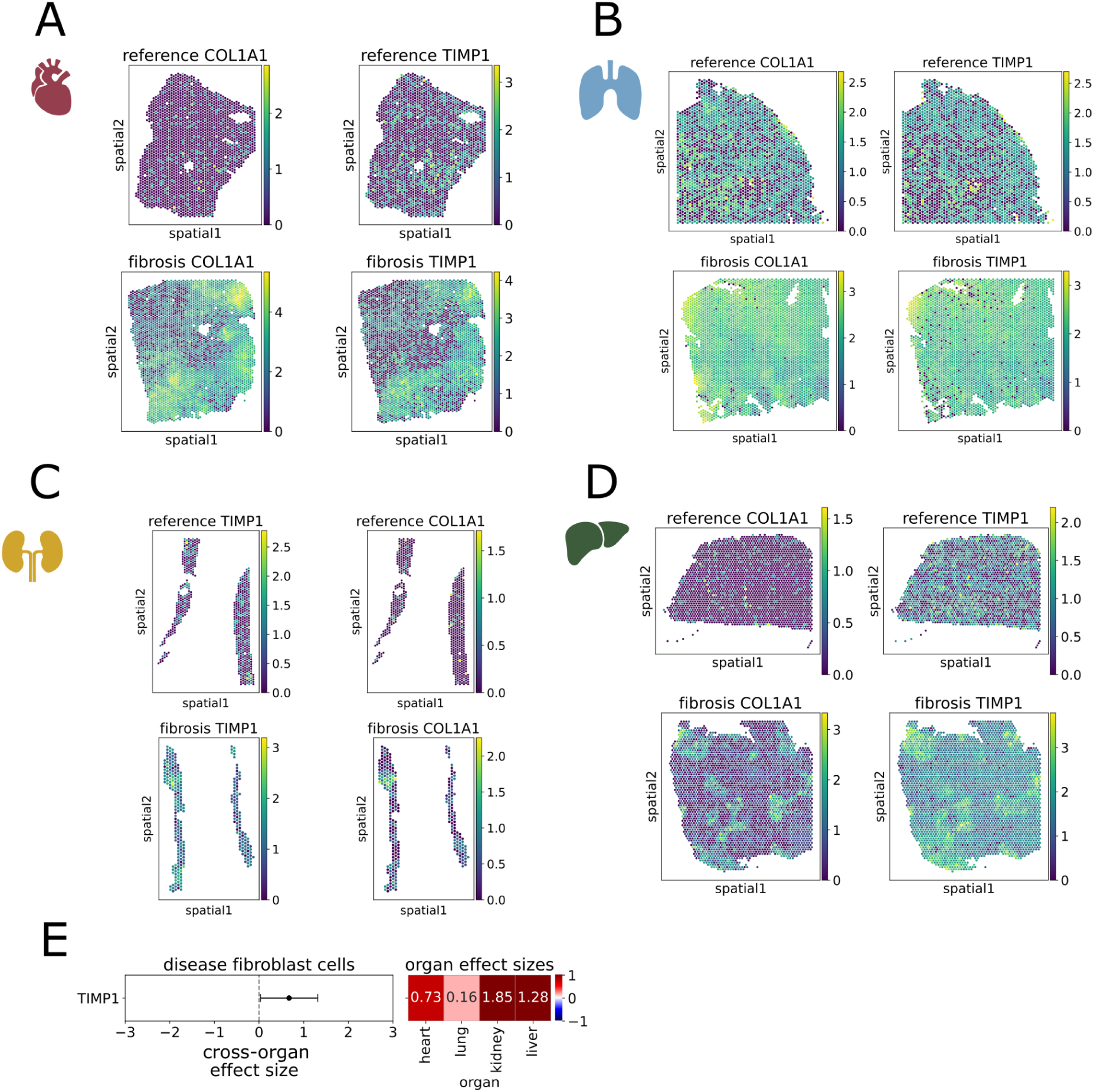
TIMP1 in fibrosis. Spatial COL1A1 (left) and TIMP1 (right) expression in one exemplary Visium spatial transcriptomics slide from a reference (top) or a fibrosis (bottom) sample in (A) heart, (B) lung, (C) kidney, and (D) liver. (E) TIMP1 gene regulation on disease associated fibroblasts per organ and across organs. Boxplot on the left displays the cross-organ effect size, whereas heatmap on the right shows the organ-effect sizes for each organ.

**Supplementary Figure 9.**
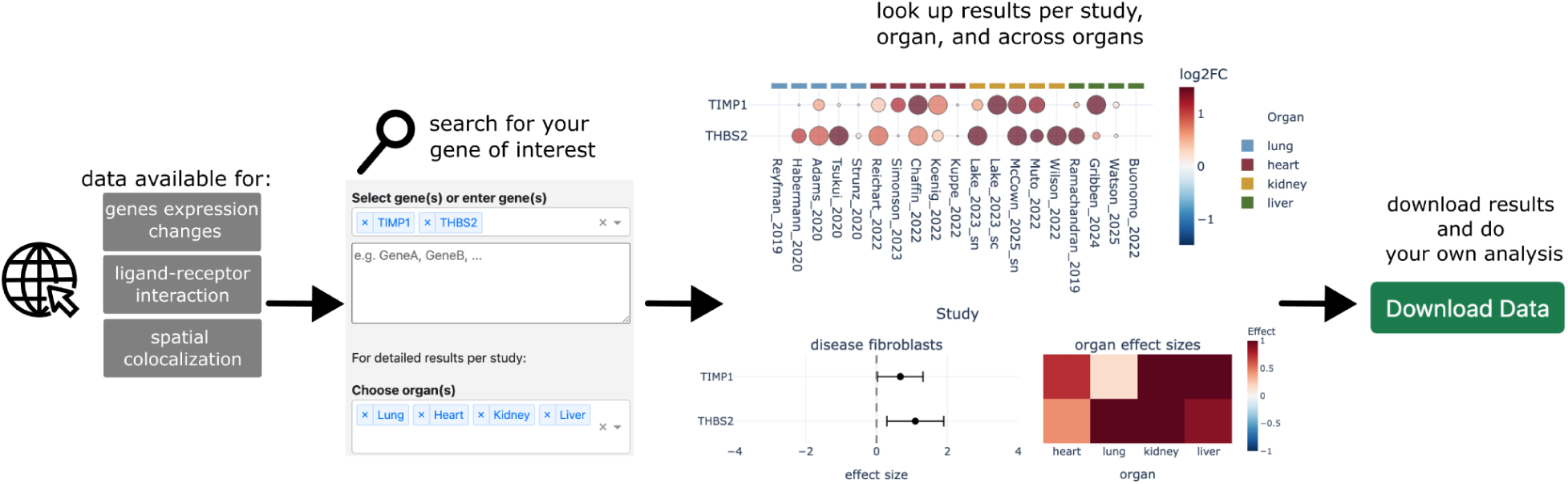
Resource website overview. Website can be reached at https://organfibrosis.saezlab.org/. Genes of interest are searchable and can be visualized. Plotted data of all available genes can be easily downloaded to enable further analyses.

